# Likelihood-free nested sampling for biochemical reaction networks

**DOI:** 10.1101/564047

**Authors:** Jan Mikelson, Mustafa Khammash

## Abstract

The development of mechanistic models of biological systems is a central part of Systems Biology. One major challenge in developing these models is the accurate inference of the model parameters. In the past years, nested sampling methods have gained an increasing amount of attention in the Systems Biology community. Some of the rather attractive features of these methods include that they are easily parallelizable and give an estimation of the variance of the final Bayesian evidence estimate from a single run. Still, the applicability of these methods is limited as they require the likelihood to be available and thus cannot be applied to stochastic systems with intractable likelihoods. In this paper, we present a likelihood-free nested sampling formulation that gives an unbiased estimator of the Bayesian evidence as well as samples from the posterior. Unlike most common nested sampling schemes we propose to use the information about the samples from the final prior volume to aid in the approximation of the Bayesian evidence and show how this allows us to formulate a lower bound on the variance of the obtained estimator. We proceed and use this lower bound to formulate a novel termination criterion for nested sampling approaches. We illustrate how our approach is applied to several realistically sized models with simulated data as well as recently published biological data. The presented method provides a viable alternative to other likelihood-free inference schemes such as Sequential Monte Carlo or Approximate Bayesian Computations methods. We also provide an intuitive and performative C++ implementation of our method.

## 1 Introduction

The accurate modelling and simulation of biological processes such as gene expression or signalling has gained a lot of interest over the last years, resulting in a large body of literature addressing various types of models along with the means for their identification and simulation. The main purpose of these models is to qualitatively or quantitatively describe observed biological dynamics while giving insights into the underlying bio-molecular mechanisms.

One important aspect in the design of these models is the determination of the model parameters. Often there exists a mechanistic model of the cellular processes, but their parameters (e.g. reaction rates or initial molecule concentrations) are largely unknown. Since the same network topology may result in different behaviour depending on the chosen parameters [26], this presents a major challenge for modelling and underscores the need for effective parameter estimation techniques.

The models used in Systems Biology can be coarsely classified into two groups: deterministic and stochastic models. Deterministic models usually rely on ordinary differential equations which, given the parameters and initial conditions, can describe the time evolution of the biological system in a deterministic manner. However, many cellular processes like gene expression are subject to random fluctuations [12, 36], which can have important biological functions [43, 49, 31] as well as contain useful information about the underlying molecular mechanisms [39]. The important role of stochastic fluctuations in biological systems has lead to increased interest in stochastic models and methods for their parameter inference[3, 25, 32, 41, 42, 56]. Such stochastic models are usually described in the framework of stochastic chemical reaction networks that can be simulated using Gillespie’s Stochastic Simulation Algorithm (SSA) [17]. In recent years, the availability of single-cell trajectory data has drastically increased, providing detailed information about the (potentially stochastic) development of single cells throughout time.

Despite the increasing interest in stochastic systems, performing inference on them is still challenging and the available methods are computationally very demanding (see for instance [3, 20, 53]). Common algorithmic approaches for such cases include various kinds of sequential Monte Carlo methods (SMC) [9, 6], Markov Chain Monte Carlo (MCMC) methods [19, 3, 45], approximate Bayesian computation (ABC) methods [54, 32, 28], iterative filtering [27] and nested sampling (NS) approaches [52, 29, 37, 15]. Furthermore, to reduce computational complexity, several of these inference methods rely on approximating the model dynamics (for instance using the diffusion approximation [18] or linear noise approximation [11]). However, these approximations may not always be justifiable (in the case of low copy numbers of the reactants for example) and might obscure crucial system behaviour. One particular problem that is common to most inference methods is the usually high dimensional parameter space. Most of the sampling-based inference techniques require the exploration of the full parameter space, which is not an easy task as the dimension of the parameter space increases. In this paper, we focus on nested sampling methods and investigate its applicability to stochastic systems. Coming originally from the cosmology community, NS (originally introduced in [52]) has gained increasing popularity and found also applications in Systems Biology (see for instance [1, 5, 10, 29, 46]). Several implementations of NS are available ([14, 22]) and in [29] the authors even provide a NS implementation specifically for a Systems Biology context. Even though the original purpose of NS was to efficiently compute the Bayesian evidence, it has more and more become a viable alternative to MCMC methods for the approximation of the posterior (see for instance [16, 24]).

There are various reasons for the interest in NS which are discussed in detail in [40, 33] and the references within. Some of the rather appealing features of NS is that it performs well for multimodal distributions [16, 22], is easily parallelizable [23, 5] and provides a natural means to compute error bars on all of its results without needing multiple runs of the algorithm [51, 40]. For a comparison of MCMC and NS see for instance [46, 40], for a discussion of other methods to compute the Bayesian evidence using MCMC see [40, 34]. Like standard MCMC methods, NS requires the availability of the likelihood *l*(*θ*) which limits its use to models that allow for the computation of the likelihood such as deterministic models and simple stochastic models. In this paper, we consider an extension to the original NS framework that, similarly to the particle MCMC method [55] and particle SMC [2], allows the use of approximated likelihoods instead of the actual likelihood to be used for NS. In the following we introduce the notation and problem formulation, in section 2 we revisit the basic NS idea and outline some of its features. Section 3 is dedicated to the likelihood-free NS formulation and in section 4 we demonstrate its performance on several chosen examples.

### 1.1 Chemical Reaction Networks

We are considering a *n*_*x*_-dimensional Markov Process *X*(*t*) depending on a *d*-dimensional parameter vector *θ*. We denote with *X*_*i*_(*t*) the *i*^th^ entryof the state vector at time *t* and with 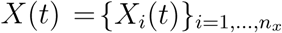 the state vector at time *t*. We will write *X*_*τ*_ = *X*(*t*_*τ*_) when talking about the state vector at a timepoint *t*_*τ*_ indexed with *τ*.

In the context of stochastic chemical reaction networks this Markov process describes the abundances of *n*_*x*_ species 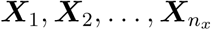, reacting through *n*_*R*_ reactions 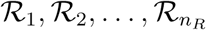 written as

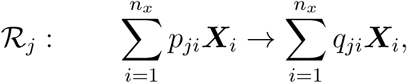

where *p*_*ji*_ is the numbers of molecules of species ***X***^*i*^ involved in reaction *R*_*j*_, and *q*_*ji*_ is the number of molecules of species ***X***^*i*^ produced by that reaction. The random variable *X*_*i*_(*t*) corresponds to the number of molecules of species ***X***^*i*^ at time *t*. Each reaction ℛ^*j*^ has an associated propensity. The reaction propensities at a given time *t* depend on the current state *X*(*t*) and on a d-dimensional parameter vector *θ*.

### 1.2 General Task

The process *X*(*t*) is usually not directly observable but can only be observed indirectly through a *n*_*y*_-dimensional observation vector

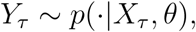

which depends on the state *X*_*τ*_ and on the *d*-dimensional parameter vector *θ* ∈ Ω where Ω ⊆ ℝ*^d^* denotes the parameter space. We shall assume that the variable *Y* is not observed at all times but only on *T* timepoints *t*_1_*, …, t_T_* and only for *M* different trajectories. With ***y*** we denote the collection of observations at all time points. In the Bayesian approach the parameter vector *θ* is treated as a random variable with associated prior *π*(*θ*). The goal is not to find just one set of parameters, but rather to compute the posterior distribution 𝒫(*θ*|***y***) of *θ*

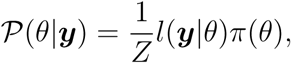

where *l*(***y***|*θ*) (we will also write *l*(*θ*) if the dependence on ***y*** is clear from the context) is the likelihood of *θ* for the particular observation ***y*** and *Z* is the Bayesian evidence

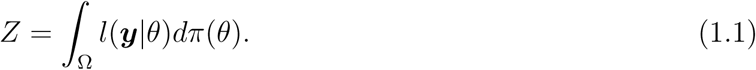

This has several advantages over a single point estimate as it gives insight into the areas of the parameter space resulting in model behaviour similar to the observations as well as about their relevance for the simulation outcome (a wide posterior indicates non-identifiability for example). For a detailed discussion of Bayesian approaches see for instance [34]. In this paper we follow the Bayesian approach and aim to recover the posterior 𝒫(*θ*|***y***). In the following section we briefly outline the basic nested sampling approach.

## 2 Nested Sampling

Nested sampling is a Bayesian inference technique that was originally introduced by John Skilling in [52] to compute the Bayesian evidence 1.1. NS can be viewed as an importance sampling technique (as for instance discussed in [47]) as it approximates the evidence by generating samples *θ*_*i*_, weights *w*_*i*_ and likelihoods *l*_*i*_ = *l*(*θ*_*i*_) such that the weighted samples (*θ*_*i*_, *w*_*i*_) can be used to obtain numerical approximations of a function *f* over the prior *π*

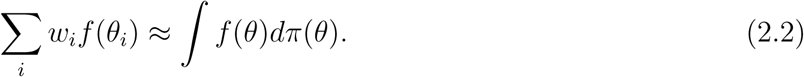

To compute an approximation 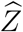 of the Bayesian evidence 1.1, *f* is chosen to be the likelihood function *l*

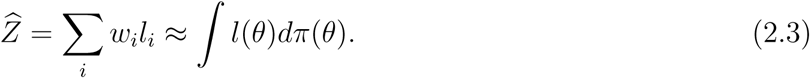

The points *θ*_*i*_ are sampled from the prior distribution constrained to super level sets of the likelihood corresponding to an increasing sequence of thresholds. In this sense it can also be viewed as a sequential Monte Carlo method, where the intermediate distributions are the nested super level sets of the likelihood. This way, samples from NS are concentrated around the higher regions of the likelihood. One can also use the weights *l*_*i*_ × *w*_*i*_ instead of *w*_*i*_ to approximate functions over the posterior 𝒫(*θ*)

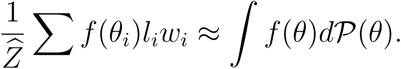

### 2.1 NS algorithm

In the following we briefly outline the NS algorithm. First, a set ℒ_0_ of *N* “live” particles {*θ*^*i*^}_*i*=1*,…,N*_ is sampled from the prior *π*

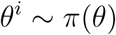

and their likelihoods *l*_*i*_ = *l*(*θ*^*i*^) are computed. Then the particle with the lowest likelihood

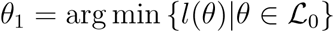

gets removed from the set of live particles and saved together with its likelihood

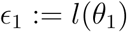

in a set of “dead” particles 𝒟. A new particle *θ*^*^ is then sampled from the prior under the constraint that its likelihood is higher than *ϵ*_1_

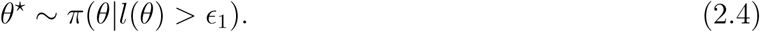

This particle is combined with the remaining particles of ℒ_0_ to form a new set of live particles ℒ_1_ that are now distributed according to the constrained prior *π*(*θ*|*l*(*θ*) > *ϵ*_1_), which we denote as

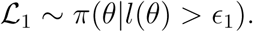

This procedure is repeated until a predefined termination criteria is satisfied. The result is a sequence of dead points *θ*_*i*_ with corresponding likelihoods *ϵ*_*i*_ that are concentrated in the regions of high likelihood. The Nested Sampling procedure is shown in Algorithm 1.

#### Algorithm 1 Nested sampling algorithm

1: Given observations ***y*** and a prior *π*(*θ*) for *θ*.

2: Sample *N* particles *θ*^*k*^ from the prior *π* and save in the set ℒ_0_, set = 𝒟{∅}

3: **for** i = 1, 2, …, m **do**

4: Set *θ*_*i*_ = arg min {*l*(*θ*)|*θ* ∈ ℒ_*i-*1_} and *ϵ*_*i*_ = *l*(*θ*_*i*_)

5: Add {*θ*_*i*_, *ϵ*_*i*_} to 𝒟

6: Set ℒ^*i*^ = ℒ_*i-*1_\*θ*_*i*_

7: Sample *θ*^*^ ∼ *π*(*θ l*(*θ*) > *ϵ*_*i*_) and add it to ℒ^*i*^

8: **end for**

### 2.2 Approximating the Bayesian Evidence

Nested sampling exploits the fact that the Bayesian evidence 1.1 can also be written^1^ (see [52]) as a one dimensional integral

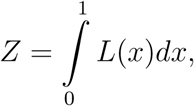

over the prior volume

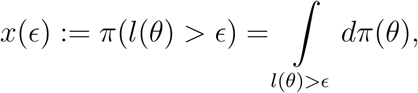

where *L*(*x*) denotes the likelihood corresponding to the constrained prior with volume *x*

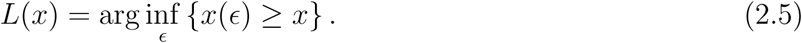

We have visualized these quantities on a simple example with a uniform prior on [0, 1] in Figure 1.

**Figure 1:**
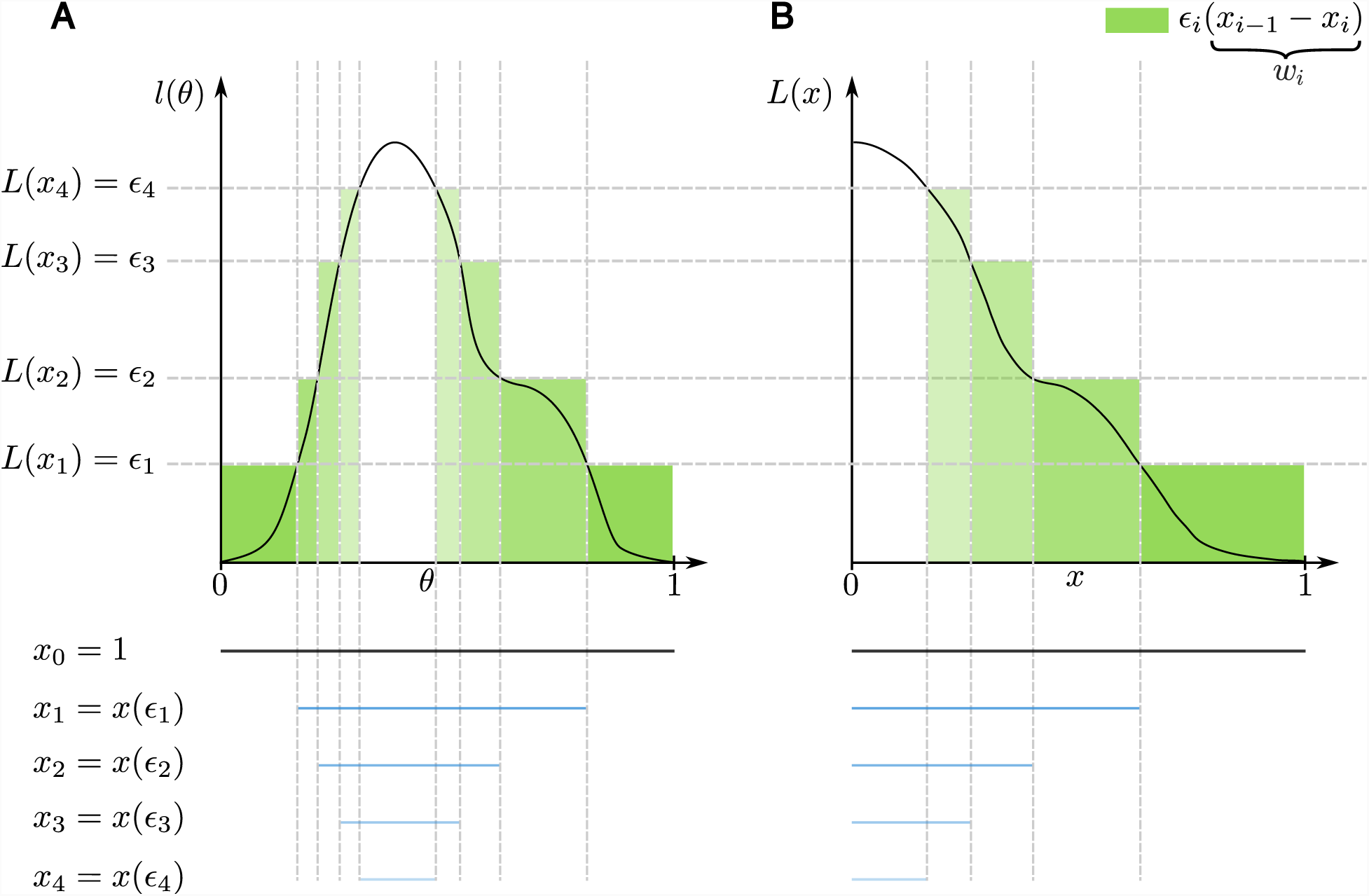
Illustration of the nested sampling approximation with a uniform prior on [0, 1]. **A:** The integral over the parameter space ∫_Ω_*l*(*θ*)*dθ*. **B:** The transformed integral 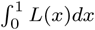 over the prior volume *x*.

The sampling scheme of nested sampling provides a sequence of likelihoods *ϵ*_1_ < *ϵ*_2_ < *…* < *ϵ*_*m*_, but their corresponding prior volumes *x*(*ϵ*_*i*_) are not known. However, since the *ϵ*_*i*_ are obtained by iteratively removing the lowest likelihood of *N* uniformly distributed points on the constrained prior *π*(*θ*|*l*(*θ*) > *ϵ*_*i-*1_), the prior volume *x*(*ϵ*_*i*_) can be written as

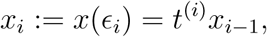

where each *t*^(*i*)^ is an independent sample of the random variable *t* which is distributed as the largest of *N* uniform random variables on the interval [0, 1] and *x*_0_ = 1 (For further justification and discussion on this see [52, 15, 8] and the references within). The values *t*^(*i*)^ are not known and need to be estimated. Since their distribution is known^2^, they can be approximated by their means 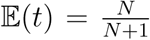 (or by the mean of their logs 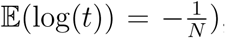), and thus the *i*^th^ prior volume can be approximated as

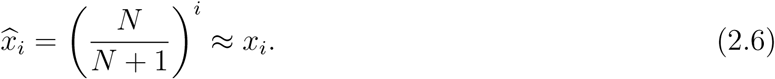

With these prior volumes one can compute the importance weights *w*_*i*_ in equation 2.2 and 2.3 for each of the dead particles *θ*_*i*_ as

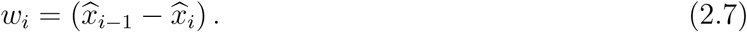

These weights correct for the fact that the samples in 𝒟 are not drawn uniformly from the prior, but are concentrated in areas of high likelihood. We note that to integrate a function on the parameter space Ω over the prior *π*, as in equations 2.2, only these weights are needed. To approximate *Z*, NS uses these weights to integrate the likelihood function *l*(*θ*) over the prior

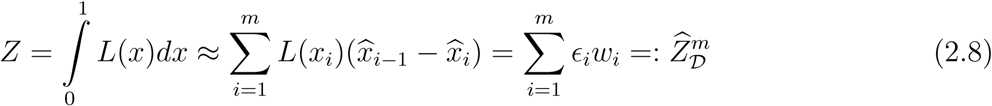

where *m* is the number of performed NS iterations and the subscript 𝒟 in 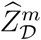 emphasizes that for NS the evidence estimate is obtained using only the dead points in 𝒟. The justification for these weights as well as an in depth discussion and error approximation can be found in [8, 24, 30] and the references therein. This basic idea of nested sampling has seen several modifications and improvements over the years, along with in-depth discussions of various sampling schemes for the constrained prior [14, 22], parallel formulations [22, 23, 5] and several implementations [14, 22, 29].

### 2.3 Termination of NS

Assuming that the distribution 2.4 can be efficiently sampled, each iteration of the NS scheme has the same computational complexity (the computationally most expensive step is usually to sample *θ*^*^ ∼ *π*(*θ*|*l*(*θ*) > *ϵ*_*i*_) and computing its likelihood). The NS algorithm is usually run until the remaining prior volume multiplied by the highest likelihood in this volume is smaller than a predefined fraction of the current BE estimate (see [52]). We write this quantity as

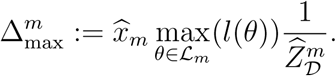

Some other termination criteria have been suggested (for instance in [22]), but since the prior volume decreases exponentially with the number of NS iterations and each iteration takes the same computational time, the choice of the particular termination criterion is not critical.

### 2.4 Parallelization of NS

The parallelization of NS can be done in a very straight forward manner. Still several different parallelization schemes have been suggested in [22, 23, 5] (for a short overview see section S1). We use a parallelization scheme similar to the one presented in [23], where at each iteration not only the one particle with the lowest likelihood is resampled, but the *r* lowest particles. The resulting parallel scheme is outlined in Algorithm 2. With *r* parallel particles the final approximation 2.8 changes to

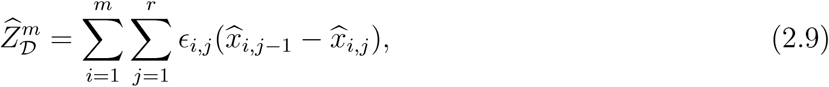

with 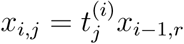 and 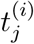 being *i*^th^ sample of *t*_*j*_ which is the *j*^th^ largest number among *N* uniform numbers between 0 and 1 ^3^ (with the obvious boundary condition *x*_0,*r*_ = 1). We note that this is slightly different than the parallelization scheme presented in [22, 23, 5], for a brief discussion see S1.

#### Algorithm 2: Parallel nested sampling algorithm. The samples drawn in line 11 are all independent and thus can be drawn in parallel

1: Given observations ***y*** and a prior *π*(*θ*) for *θ*.

2: Sample *N* particles *θ*^*k*^ from the prior *π* and save them in the set ℒ_0_, set 𝒟 = {∅}

3: **for** i = 1, 2, …, m **do**

4: **for** j = 1, 2, …, r **do**

5: Set *θ*_*i,j*_ = arg min {*l*(*θ*)*|θ ∈* ℒ_*i-*1_} and *ϵ*_*i,j*_ = *l*(*θ*_*i,j*_)

6: Add {*θ*_*i,j*_, *ϵ*_*i,j*_} to 𝒟

7: remove *θ*_*i,j*_ from ℒ_*i-*1_

8: **end for**

9: Set ℒ^*i*^ = ℒ_*i-*1_

10: **for** j = 1, 2, …, r **do**

11: Sample *θ*^*^ *∼ π*(*θ | l*(*θ*) *>, ϵ_i,r_*) and add it to ℒ^*i*^

12: **end for**

13: **end for**

## 3 Likelihood-free nested sampling (LF-NS)

In many cases (such as most of the above mentioned stochastic models) the likelihood *l*(*θ*) cannot be directly computed, making approaches like MCMC methods or nested sampling not applicable. Fortunately, many variations of MCMC have been described circumventing this problem by introducing likelihood-free MCMC methods such as [35] or [55] as well as other likelihood-free methods such as ABC [54] or likelihood-free sequential Monte Carlo (SMC) methods [50]. These approaches usually rely on forward simulation of a given parameter vector *θ* to obtain a simulated data set that can then be compared to the real data or can be used to compute a likelihood approximation 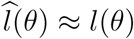. In the following we briefly illustrate one way to approximate the likelihood.

### 3.1 Likelihood approximation using particle filters

A common way to approximate the likelihood through forward simulation is using a particle filter (see for instance [44] or [19]), which iteratively simulates the stochastic system with *H* particles and then resamples these particles. In the following we illustrate such a particle filter likelihood approximation on a simple birth death model, where one species (mRNA) is produced at rate *k* = 1 and degrades at rate *γ* = 0.1. We simulated one trajectory (shown in Figure 2 A) of this system using Gillespies stochastic simulation algorithm (SSA [17]) and, using the finite state projection (FSP [38]), computed the likelihood *l*(*k*) for different values of *k* while keeping *γ* fixed to 0 .1. The true likelihood for different *k* is shown as the solid red line in Figure 2 B and C. We also illustrated the likelihood approximation 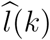 using a particle filter ([19]) with *H* = 100 particles for three values of *k*. For each of the values for *k* we computed 1000 realizations of 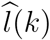 and plotted the empirical distributions in Figure 2 C. Note that 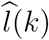 is itself a random variable with distribution 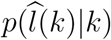 and has a mean equal to the true likelihood 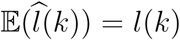 (see for instance [44]). We also sampled 10^6^ values of *k* from a log uniform prior and approximate for each *k* its likelihood with the same particle filter with *H* = 100 particles. We plotted the contour lines of this joint distribution in Figure 2 B.

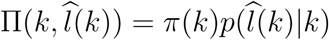

In the following we discuss how to utilize such a likelihood approximation to apply the above described NS procedure to cases where the likelihood is not available. Throughout the paper we assume that the likelihood approximation 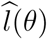 is obtained using a particle filter, but our result hold for any unbiased likelihood estimator.

**Figure 2:**
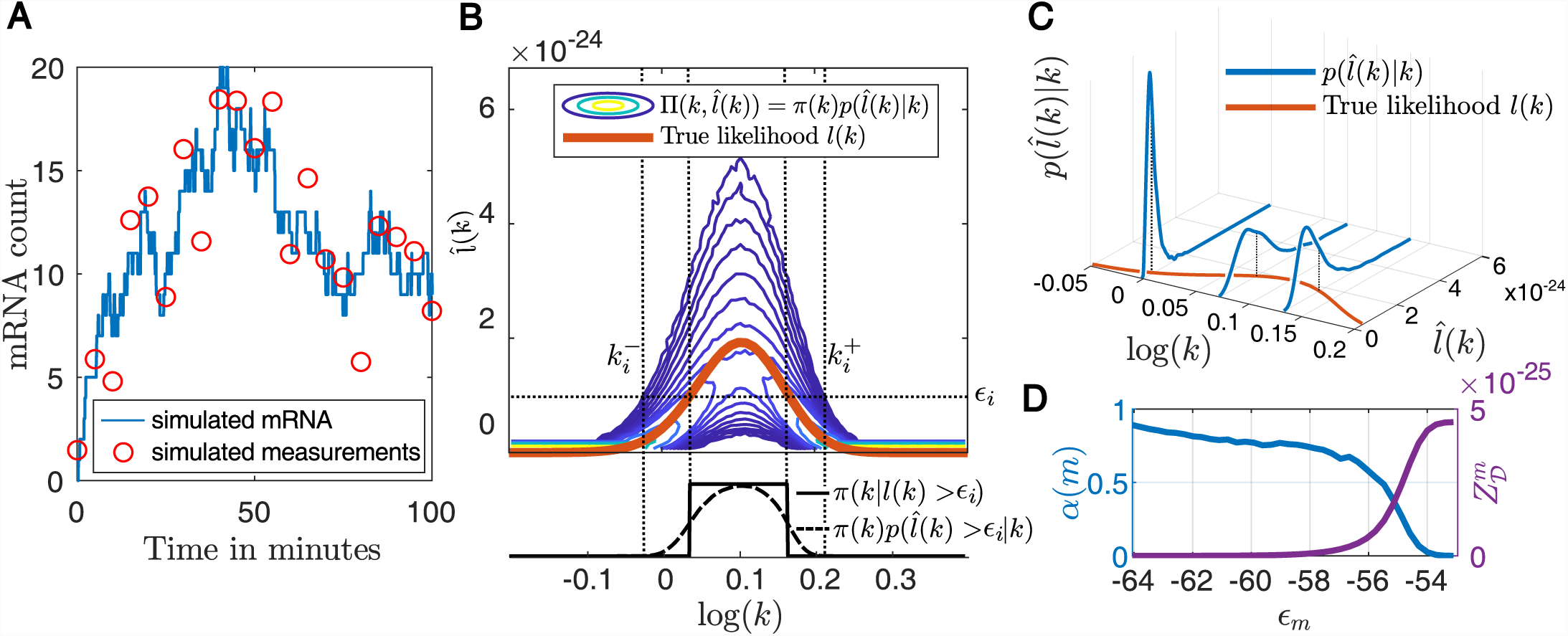
**A:** A simulated trajectory of the birth death system using k = 1 and *γ* = 0.1 with 21 equally spaced measurements (taken to be normally distributed around the mRNA count with *σ* = 2).**B:** Top: Likelihood for different parameters k (red) and contour lines of the joint distribution Π(*k,* log(*k*) of the parameter *k* and its likelihood approximation 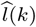, based on 10^6^ samples of the likelihood approximation obtained with a particle filter with 100 particles. Bottom: The constrained priors 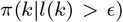 and 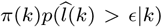 for *ϵ* = 1*e -* 24. **C:** Example distributions 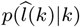 (blue) for *k* = 1, 1.2 and 1.4 and the true likelihood *l*(*k*) red. **D:** blue: The ratio *α*(*m*) of the probability masses of 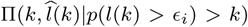 above and below each likelihood threshold *ϵ*_*i*_, constrained to those regions of *k* with 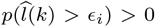 (these are the parameter regions in panel B between 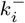 and 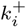). purple: The evidence as estimated by all particles with likelihood below 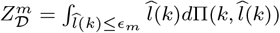.

### 3.2 The LF-NS scheme

From here on we assume that the true likelihood *l*(*θ*) is not available, but a realization 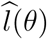 of the approximated likelihood having the distribution

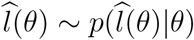

with

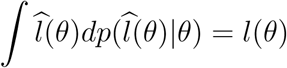

can be computed.

For NS, the constraint prior *π*(*θ|l*(*θ*) *> ϵ_i_*) needs to be sampled. Since in the likelihood-free case, the likelihood *l*(*θ*) is not available and 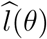 is itself a random variable, the set 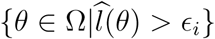 (which is the support of the constrained prior) is not defined. To apply the NS idea to the likelihood-free case, we propose to perform the NS procedure on the joint prior

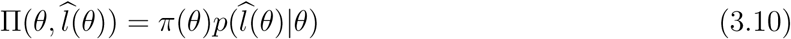

on the set Ω *×* ℝ_*>*0_. This joint prior can be sampled by drawing a sample *θ*^*\**^ from the prior *π*(*θ*) and then drawing one sample 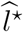 from the distribution of likelihood approximations 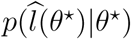. With such a sampling scheme we perform the NS steps of constructing the set of “dead” particles 𝒟 on the joint prior 3.10. As in standard NS, we sample a set of *N* “live” particles 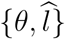 from 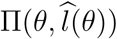, then we iteratively remove the particle 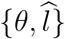 with the lowest likelihood sample 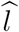 from the set of live points and add it to the dead points. The LF-NS algorithm is shown in Algorithm 3.

The parallel version of LF-NS is analogous to the parallelization of the standard NS algorithm in Algorithm 2.

#### Algorithm 3: Likelihood-free nested sampling algorithm

1: Given observations ***y***, a prior *π*(*θ*) for *θ* and a likelihood approximation 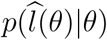.

2: Sample *N* particles 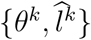 from the prior 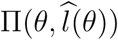 and save it in the set ℒ_0_, set 𝒟 = {*∅*}

3: **for** i = 1, 2, …, m **do**

4: Find 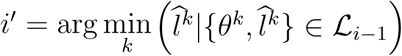 and set 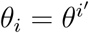 and 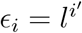

5: Add {*θ*_*i*_, *ϵ*_*i*_} to 𝒟

6: Set ℒ^*i*^ = ℒ_*i-*1_\{*θ*_*i*_, *ϵ*_*i*_}

7: Sample 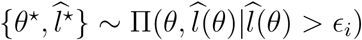 and add it to ℒ_*i*_

8: **end for**

### 3.3 LF-NS is unbiased

As for standard NS, the sampling procedure for LF-NS guarantees that each set of live points ℒ^*i*^ contains *N* samples uniformly distributed according to the constrained joint prior 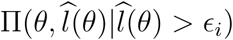, thus removing the sample with the lowest likelihood approximation 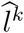 results in the same shrinkage of prior volume as the standard NS scheme. The prior volumes *x*_*i*_ = *t*^(*i*)^*x*_*i-*1_ now correspond to the volumes of the constraint joint priors 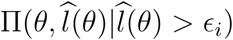 and the resulting weights 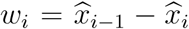 can be used, similarly as in equation 2.2, to integrate functions *f* over the constrained prior

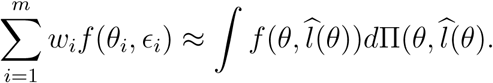

Using *f* (*θ*_*i*_, *ϵ*_*i*_) = *ϵ*_*i*_ we can use this to approximate the Bayesian Evidence

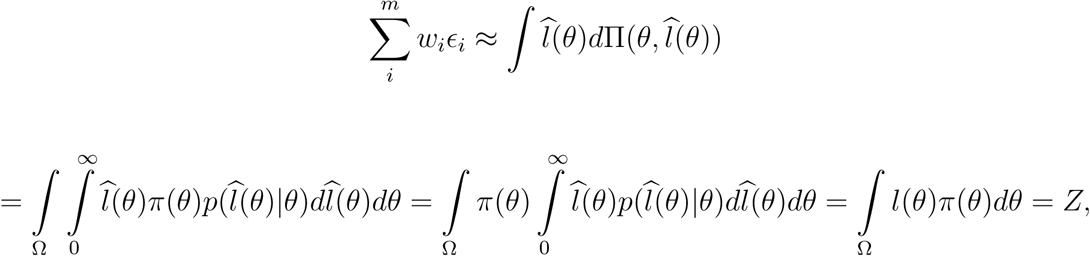

where the last equality relies on the unbiasedness of 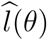.

While the procedure for LF-NS is very similar to the standard NS algorithm, the new samples *θ*^*\**^ have to be drawn from the constraint joint prior 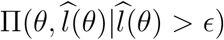 instead from the constrained prior *π*(*θ|l*(*θ*) *> ϵ*). In the following we discuss the resulting difficulties and show how to overcome them.

### 3.4 Sampling from the super-level sets of the likelihood

One of the main challenges [7, 40, 4] in the classical NS algorithm is the sampling from the prior constrained to higher likelihood regions *π*(*θ|l*(*θ*) *> ϵ*). A lot of effort has been dedicated to find ways to sample from the constraine prior efficiently, the most popular approaches include slice sampling [22] and ellipsoid based sampling [16].

In the case of LF-NS, at the *i*^th^ iteration we are sampling not just a new parameter vector *θ*^*\**^ but also a realization of its likelihood approximation 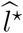 from

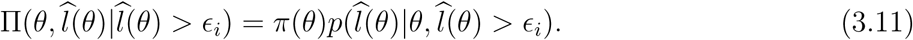

Since it is in general not possible to sample 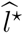 from the constraint distribution 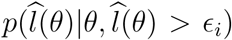 directly, we sample *θ*^*\**^ from the prior *π*(*θ*), then sample 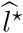 from the unconstrained distribution 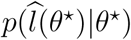 and accept the pair 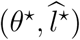 only if 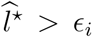. While this procedure guarantees that the resulting samples are drawn from 3.11, the acceptance rate might become very low. Each live set ℒ^*i*^ consists of *N* pairs 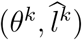 distributed according to 3.11, thus the parameter vectors *θ*^*k*^ in ℒ^*i*^ are distributed according to

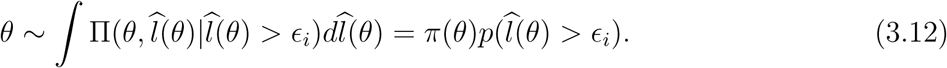

We plotted an example of the distributions 2.4 and 3.12 for the example of the birth-death process in Figure 2 B. The distribution 3.12 has usually an infinite support, although in practice 3.12 will be close to zero for large areas of the parameter space Ω. Similarly to NS, we propose to use the set ℒ^*i*^ to drawn from the areas where 3.12 is non zero. Slice sampling methods ([1, 22]) are unfortunately not applicable for LF-NS since they require a way to evaluate its target distribution at each of its samples. We can still use ellipsoid sampling schemes, but unlike in the case of NS where the target distribution *π*(*θ|l*(*θ*) *> ϵ*) has compact support, the target distribution for LF-NS 3.12 has potentially infinite support framing ellipsoid based sampling approaches rather unfitting. Sampling using MCMC methods (as suggsted in [52]) is expected to work even for target distributions with infinite support, but suffer from the known MCMC drawbacks, as they produce correlated samples and might get stuck in disconnected regions.

To account for the smooth shape of 3.12 we propose to employ a density estimation approach. At each iteration *i*, we estimate the density 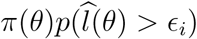 from the live points and employ a rejection sampling approach to sample uniformly from the prior on the domain of this approximation. As density estimation technique, we use Dirichlet Process Gaussian Mixture Model (DP-GMM) [21], which approximates the distribution 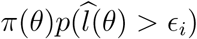 with a mixture of Gaussians. DP-GMM uses a hierarchical prior on the mixture model and assumes that the mixture components are distributed according to a Dirichlet Process. The inference of the distribution is an iterative process that uses Gibbs sampling to infer the number and shape of the Gaussians as well as the parameters and hyper parameters of the mixture model. DP-GMM estimations perform comparably well with sparse and high dimensional data and are less sensitive to outliers. Further, since we employ a parallelized LF- NS scheme, the density estimation has to be performed only after the finish of each parallel iteration, making the computational effort of density estimations negligible compared to the computational effort for the likelihood approximation. For a detailed comparison between DP-GMM and kernel density estimation and a further discussion of DP-GMM see [21], for an illustration of DP-GMM, KDE and ellipsoid sampler see section S2. Even though for the presented examples we employ DP-GMM, we note that in theory any sampling scheme that samples uniformly from the prior *π*(*θ*) over the support of 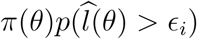 will work.

### 3.5 A lower bound on the estimator variance

Unlike for NS, for LF-NS, even if at each iteration the proposal particle *θ*^*\**^ is sampled from the support of 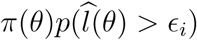, it will only be accepted with probability 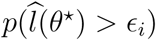. This means that depending on the variance of the likelihood estimation 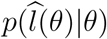 and the current likelihood threshold *ϵ*_*i*_ the acceptance rate for LF-NS will change and with it the computational cost. We illustrated this on the example for the birth-death model above. For each of the 10^6^ samples 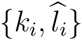 from 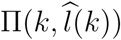 we set *ϵ*_*i*_ = *l*_*i*_ and considered the particles 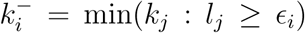 and *k*^+^ = max(*k*_*j*_ : *l*_*j*_ *≥ ϵ*_*i*_) (illustrated in Figure 2 B). The particles {*k*_*j*_} in between 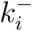 and 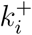 give a numerical approximation of the support of 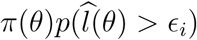. We denote with 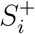 all the pairs 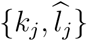 with *k*_*j*_ between 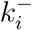 and 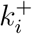 with a likelihood above *ϵ*_*i*_ and with 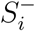 the pairs with a likelihood below *ϵ*_*i*_

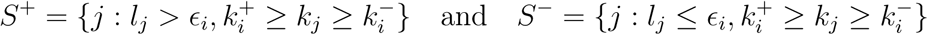

and computed the ratio of the number of their element

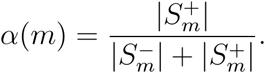

The values of *α*(*m*) give us an idea what the acceptance rate for LF-NS looks like in the best case where the new particles *k*^*\**^ are sampled from the support of 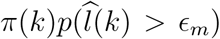. We plotted *α*(*m*) in Figure 2 D as well as the evidence 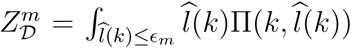. We see that *α*(*m*) decreases to almost zero as 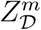 approaches *Z*. The shape of *α*_*m*_ will in general be dependent on the variance of the likelihood approximation 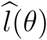. For a further discussion on the acceptance rate for different particle filter settings see section S3.

Due to this possible increase in computational time, it is important to terminate the LF-NS algorithm as soon as possible. We propose to use for the Bayesian evidence estimation not only the dead particles 𝒟, but also the current live points ℒ_*m*_. This possibility has been already mentioned in other places (for instance in [8, 24, 30]) but is usually not applied, since the contribution of the live particles decreases exponentially with the number of iterations.^4^ Since for standard NS each iteration is expected to take the same amount of time, most approaches simply increase the number of iterations to make the contribution of the live particles negligibly small.

The Bayesian evidence can be decomposed as

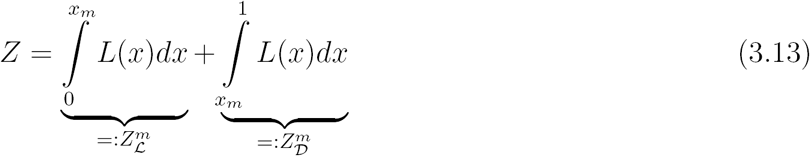

where *x*_*m*_ is the prior volume for iteration *m*. The first integral 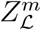 imated through the *N* live samples at any given iteration, while the integral 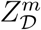 through the dead samples. Writing

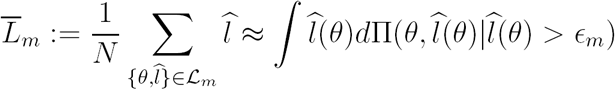

for the estimator of the integral of the likelihoods in the live set, we propose the following estimator for *Z*

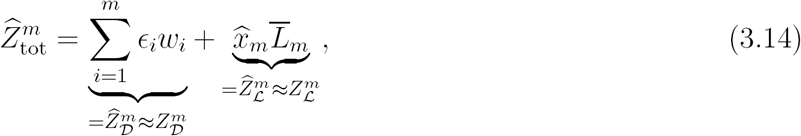

where 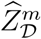 approximates the finite sum 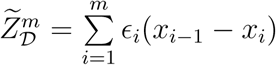 by replacing the random variables *x*_*i*_ with their means 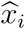 Since 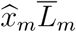 is an unbiased estimator of 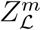 and 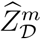is an unbiased estimator of 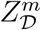, the estimator 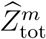 is an unbiased estimator of the Bayesian evidence *Z* for any *m*. In particular, this implies that terminating the LF-NS algorithm at any iteration *m* will result in an unbiased estimate for *Z*. However, terminating the LF-NS algorithm early on will still result in a very high variance of the estimator. Since the error of replacing the integral *Z* with the finite sum 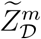 is negligible compared to the error resulting from replacing *x*_*i*_ with 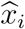 (see [13] or [8]), this variance is a result of the variances in *x*_*i*_ and the variance in the Monte Carlo estimate 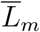.^5^ In the following we formulate a lower bound 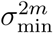 on the estimator variance 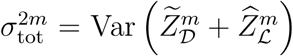 at iteration *m*, show that this lower bound is monotonically increasing in *m* and propose to terminate the LF-NS algorithm as soon as the current estimator variance differs from this lower bound by no more than a predefined threshold *δ*.

Treating the prior volumes *x*_*i*_ and the Monte Carlo estimate 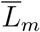 as random variables, the variance 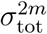 of the NS estimator at iteration *m* can be estimated at each iteration without additional computational effort (see section S4 and [30]). This variance depends on the variance of the Monte Carlo estimate 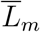 and is monotonically increasing in 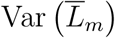 (see section S5). We define the term 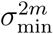 which is the same variance 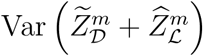 but under the additional assumption that the Monte Carlo estimate has variance 0: 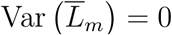. Clearly we have for any *m* (see section S5)

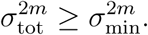

More importantly, as we show in section S5.2, 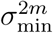 is monotonically increasing in *m*

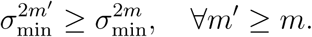

This allows us to bound the lowest achievable estimator variance 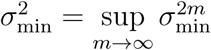 from below

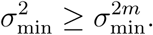

The terms for 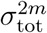 and 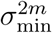 both contain the unknown value *L*_*m*_ which can be approximated using its Monte Carlo estimate 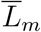 giving us the estimations of the above variances 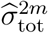 and 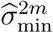. We use these variance estimates to formulate a termination criteria by defining

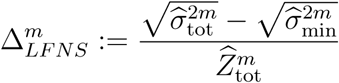

and terminate the algorithm as soon as 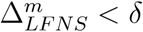 for some predefined *δ*. This termination criteria seems intuitive since it terminates the LF-NS algorithm as soon as a continuation of the algorithm is not expected to make the final estimator significantly more accurate. As a final remark we note that the final estimator 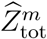 as well as the termination criteria using 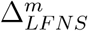 can of course also be applied in the standard NS case.

## 4 Examples

We test our proposed LF-NS algorithm on three examples for stochastic reaction kinetic models. The first example is the birth death model, already introduced in section 3.1, the second example is the Lac-Gfp model used for benchmarking in [32] and the third example is a transcriptional model from [48] with corresponding real data. In the following examples all priors are chosen as uniform or log-uniform in the bounds indicated in the posterior plots.

### 4.1 The stochastic birth-death Model

We first revisit the example of section 3.1 to compare our inference results to the solution obtained by FSP. We use the same data as in section 3.1 and use the same log-uniform prior. We run our LF-NS algorithm as described above using DP-GMM for the sampling. We used *N* = 100 LF-NS particles, *H* = 100 particle filter particles and sample at each iteration *r* = 10 particles. We ran the LF-NS algorithm until 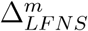 is smaller than 0.001. We show the obtained posterior in Figure 3 A. Figure 3 B shows the obtained estimates of the Bayesian evidence, where the shaded areas indicate the standard error at each iteration. The dashed red line indicates the true BE computed from 10^6^ samples from 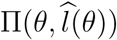. The estimates of the lower 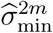 and upper bound 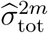 for the lowest achievable estimator variance 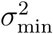 are shown in Figure 3 C and we can clearly see how they converge to the same value. For our termination criteria we show the quantities 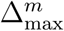 and 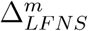 in Figure 3 D.

**Figure 3:**
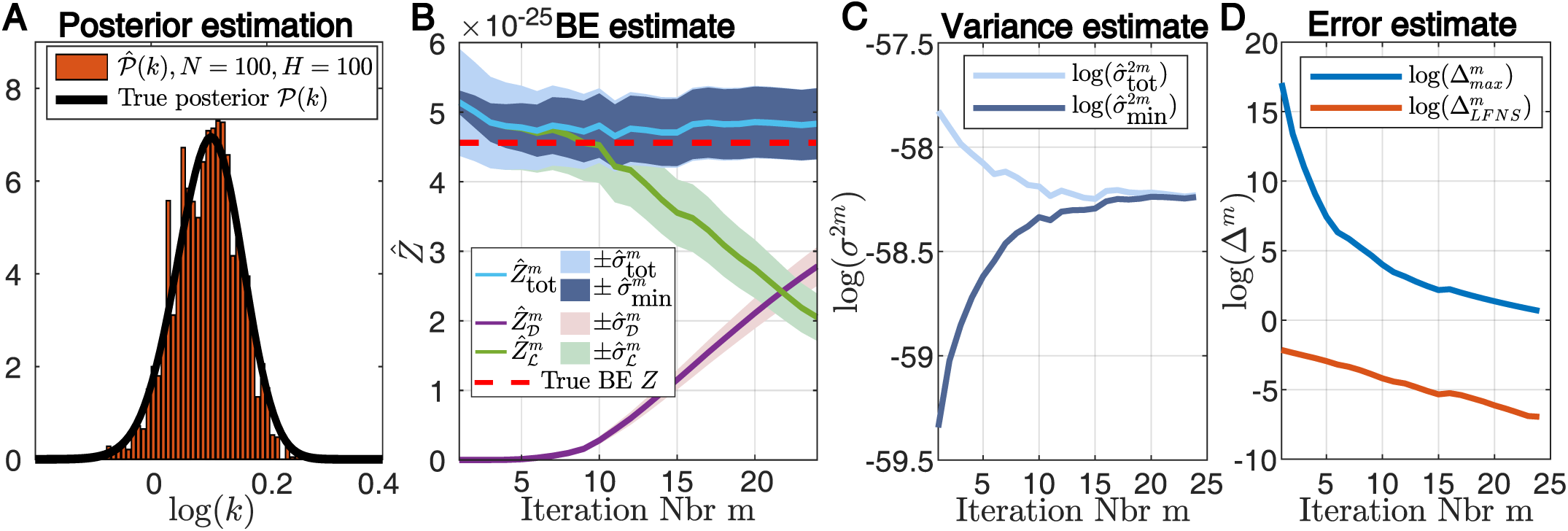
**A**: Histogram of the posterior 𝒫(*k*) estimate obtained with LF-NS using *N* = 100 and *H* = 100. The true posterior is indicated in black. **B**: Development of the estimation of the Bayesian evidence using the estimation based solely on the dead points 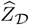, the estimate approximation from the live points 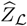 and the estimation based on both 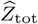. The corresponding standard errors are indicated as the shaded areas. The true Bayesian evidence is indicated with the dashed red line. **C**: Estimate of the current variance estimate 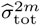 and the lower bounds for the lowest achievable variance 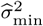. **D**: Developments of the different error estimations for each iteration.

### 4.2 The Lac-Gfp model

We demonstrate how our algorithm deals with a realistic sized stochastic model, by inferring the posterior for the parameters of the Lac-Gfp model illustrated in Figure 4 A. This model has been already used in [32] as a benchmark, although with distribution-data. Here we use the model to simulate a number of trajectories and illustrate how our approach infers the posterior of the used parameters. This model is particularly challenging in two aspects. First, the number of parameters is 18, making it a fairly large model to infer. Secondly, the model exhibits switch-like behaviour which makes it very hard to approximate the likelihood of such a switching trajectory (see section S6.2 and particularly Figure S 3 for further details). We used *N* = 500 LF-NS particles, *H* = 500 particle filter particles and sample at each iteration *r* = 50 particles.

**Figure 4:**
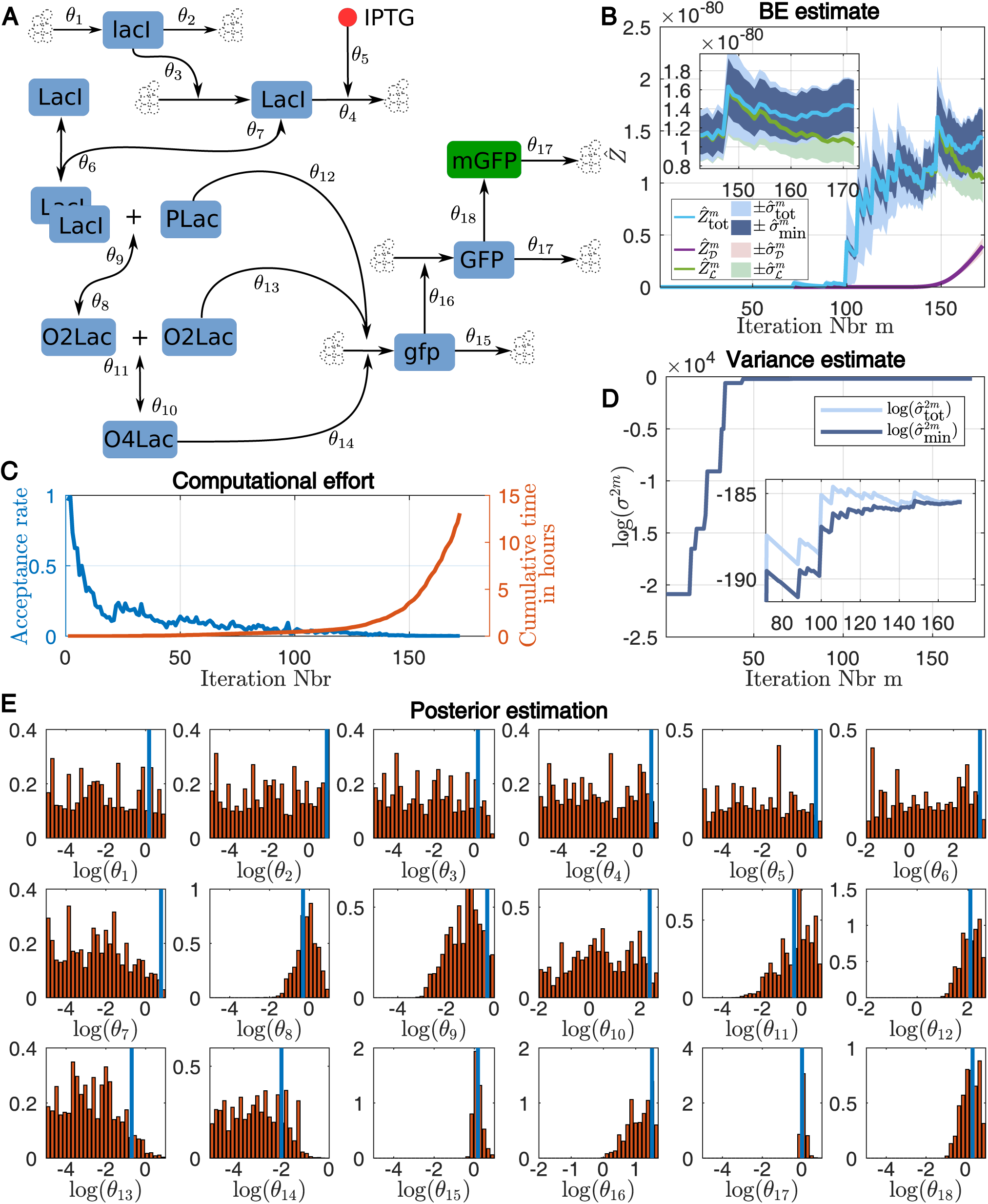
**A**: Schematic of the Lac-Gfp Model where the final measurement is the mature GFP (mGFP) and the input is IPTG (assumed to be constant 10*μ*M). **B**: Development of the estimation of the Bayesian evidence using the estimation based solely on the dead points 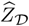, the estimate approximation from the live points 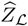 and the estimation that uses both 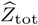. The corresponding standard errors are indicated as the shaded areas. **C**: The acceptance rate of the LF-NS algorithm for each iteration (blue) and the cumulative time needed for each iteration in hours (red). The computation was performed on 48 cores in parallel on the Euler cluster of the ETH Zurich. **D**: Estimate of the current variance estimate 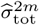 and the lower bounds for the lowest achievable variance 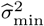. **E:** Marginals of the inferred posterior distributions of the parameters based on one simulated trajectory. The blue lines indicate the parameters used for the simulation of the data.

The measured species in this example is fluorescent Gfp (mGFP) where it is assumed that each Gfp-molecule emits fluorescence according to a normal distribution. We used one trajectory to infer the posterior, whose marginals are shown in Figure 4 E. The solid blue lines indicate the parameters used to simulate the data. Figure 4 B shows the estimated Bayesian evidence with corresponding standard errors for each iteration. Figure 4 D shows the corresponding estimations of the bounds of the lowest achievable variance. As we see, the estimated Bayesian evidence, as well as the estimated variance bounds, do several jumps in the process of the LF-NS run. These jumps correspond to iterations in which previously unsampled areas of the parameter space got sampled with a new maximal likelihood. In Figure 4 C we plotted the acceptance rate of the LF-NS algorithm for each iteration as well as the cumulative computational time^6^. The inference for this model took well over 12 hours and as we see, the computational time for each iteration seems to increases exponentially, as the acceptance rate decreases. The low acceptance rate is expected, since the number of particle filter particles *H* = 500 results in a very high variance of the particle filter estimate (see Figure S3 B). Clearly, for this example, the early termination of LF-NS is essential to obtain a solution within a reasonable time.

### 4.3 A stochastic transcription model

As a third example we use a transcription model recently used in [48], where an optogenetically inducible transcription system is used to obtain live readouts of nascent RNA counts. The model consists of a gene that can take two configurations “on” and “off”. In the “on” configuration mRNA is transcribed from this gene and can be individually measured during this transcription process (see [48] for details). We modelled the transcription through *n* = 8 subsequent RNA species that change from one to the next at a rate *λ*. This is done to account for the observed time of 2 minutes that one transcription event takes. With such a parametrization the mean time to move from species RNA_1_ to the degradation of RNA_*n*_ is 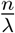. An illustration of the model is shown in Figure 5 A. For the inference of the model parameters we chose five trajectories of real biological data, shown in Figure 5 C. Clearly, the system is inherently stochastic and requires corresponding inference methods. We ran the LF-NS algorithm for *N* = 500 and *H* = 500 on these five example trajectories. The resulting marginal posteriors are shown in Figure 5 B, we also indicated the model ranges considered in [48]. These ranges were chosen in [48] in an ad-hoc manner but, apart from the values for *k*_off_ seem to fit very well with our inferred results. In Figure 5 D and E we show the development of the evidence approximation as well as the corresponding standard errors and the development of the upper and lower bound estimation for the lowest achievable variance 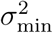. Similarly to the Lac-Gfp example, we see that the development of the BE estimate is governed by random spikes which again are due to the sampling of particles with a new highest likelihood.

**Figure 5:**
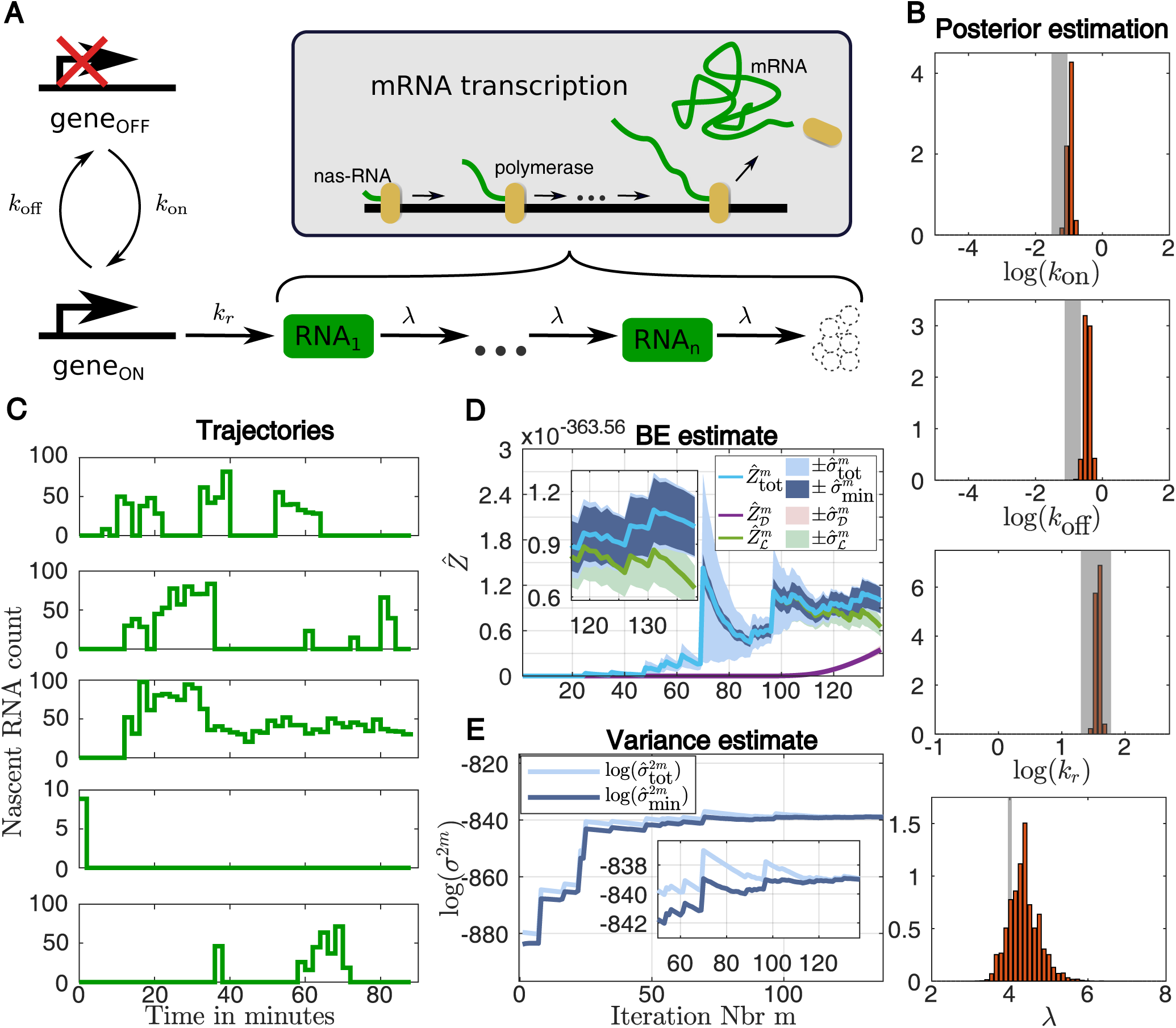
**A**: A schematic representation of the gene expression model. The model consists of a gene that switches between an “on” and an “off” state with rates *k*_on_ and *k*_off_. When “on” the gene is getting transcribed at rate *k*_*r*_. The transcription process is modelled through *n* RNA species that sequentially transform from one to the next at rate *λ*. The observed species are all of the intermediate *RNA*_*i*_ species. **B**: The marginal posterior distribution of the parameters of the system. The shaded areas indicate the parameter ranges that were considered in [48]. **C**: The five trajectories used for the parameter inference. **D**: Development of the estimation of the Bayesian evidence using the estimation based solely on the dead points 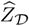, the estimate approximation from the live points 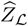 and the estimation that uses both 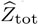. The corresponding standard errors are indicated as the shaded areas. **E**: Estimate of the current variance estimate 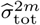 and the lower bounds for the lowest achievable variance 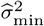.

## 5 Discussion

We have introduced a likelihood-free formulation of the well known nested sampling algorithm and have shown that it is unbiased for any unbiased likelihood estimator. While the utilization of NS for systems without an available likelihood is straight forward, one has to take precautions to avoid infeasibly high computational times. Unlike for standard NS it is crucial to include the estimation of the live samples to the final BE estimation as well as terminate the algorithm as soon as possible. We have shown how using a Monte Carlo estimate over the live points not only results in an unbiased estimator of the Bayesian evidence *Z*, but also allows us to derive a formulation for a lower bound on the achievable variance in each iteration. This lower bound at each iteration has been shown to be a lower bound for the best achievable variance and has allowed us to formulate a novel termination criterion that stops the algorithm as soon as a continuation can at best result in an insignificant improvement in accuracy. While the formulation of the variances and its lower bound were derived having a parallel LF-NS scheme in mind, they can equally well be used in the standard NS case and can be added effortlessly to already available toolboxes such as [14] or [22]. We emphasize that the lower variance bound approximation 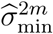 is neither a strict error term, as it only gives information of the variance of the estimator, nor a strict lower bound of the estimator variance since it contains the unknown term *L*_*m*_. Instead, it gives an estimate of the lowest achievable estimator variance that depends on the Monte Carlo estimate of the likelihood average over the live points 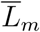. This can be seen Figure 4 D and Figure 5 E, where the lower bound estimate 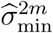 does not only make jumps, but also decreases after each jump (the actual lower bound estimate 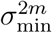 is monotonically increasing in *m* as shown in section S5.2). Our suggested LF-NS scheme has three different parameters that govern the algorithm behaviour. The number of LF-NS particles *N* determines how low the minimal variance of the estimator can get, where low values for *N* result in a rather high variance and high values for *N* result in a lower variance. The number of particles for the particle filter *H* determines how wide or narrow the likelihood estimation is and thus determines the development of the acceptance rate of the LF-NS run, while the number of LF-NS iterations determines how close the variance of the final estimate comes to the minimal variance. We have demonstrated the applicability of our method on several models with simulated as well as real biological data. Our LF-NS can, similarly to ABC, pMCMC or SMC models deal with stochastic models with intractable likelihoods and has all of the advantages of classic NS. We believe that particularly the variance estimation that can be performed from a single LF-NS run proves to be useful as well as the straight forward parallelization.

## Footnotes

1 for this to hold some weak conditions have to be satisfied, see for details [8] and [15]

2 *t* ∼ ℬ(*N,* 1) with ℬ(*a, b*) being the Beta distribution with parameters *a* and *b*.

3 This means *t*_*j*_ ∼ ℬ(*N - j* + 1*, j*)

4 We point out that while in classical nested sampling the contribution of the live points can indeed be made arbitrarily small, the resulting estimator (employing only the dead points) is strictly speaking not unbiased since it approximates the Bayesian evidence not over the full prior volume but only up to the final *x*_*m*_, which is the quantity 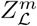 in equation 3.13

5 As pointed out in [24], when using nested sampling approximations to approximate the integral of arbitrary functions *f* over the posterior, an additional error is introduced by approximating the average value of *f* (*θ*) on the contour line of *l*(*θ*) = *ϵ*_*i*_ with the value *f* (*θ*_*i*_).

6 The computation was performed on 48 cores of the Euler cluster of the ETH Zurich.

