## Supplementary for "Likelihood-free nested sampling for biochemical reaction networks"

### S1 Parallelization schemes

#### S1.1 Parallelization in [2, 6, 5]

We briefly discuss parallelization schemes for NS as they have been discussed in several other placeses like [2, 6, 5] to better illustrate the difference between our suggested scheme.

In [5] and [2] the parallel algorithm is in principle run like the sequential version, only that at each iteration $i$ not one particle is sampled from $\pi(\theta|l(\theta) > \epsilon_i)$ but $r$. The key observation is that any particle $\theta^\star$ sampled from $\pi(\theta|l(\theta) > \epsilon_i)$ can also be accepted at iteration $j > i$ if $l(\theta^\star) > \epsilon_j$. Thus at each iteration $i$ any particle for which a likelihood gets computed, beyond the first accepted particle, is used in a subsequent iteration if its likelihood is high enough.

While this provides an intuitive parallelization of the process, the $r$ particle at each iteration $i$ are sampled from $\pi(\theta|l(\theta) > \epsilon_i)$ but are accepted only if their respective likelihoods are higher than a sequence of increasing thresholds $\epsilon_i < \epsilon_{i+1} < \ldots < \epsilon_{i+r-1}$, which may result in discarding already sampled particles. Even despite this theoretical drawback, this method works in practice very well as demonstrated in [5]. However it seems to us wasteful to potetially discard already sample particles.

In [6] the authors suggest to parallelize NS by removing $r$ particles with the lowest $r$ likelihoods from the live set at each iteration $i$ rather than just one particle. The new threshold $\epsilon_i$ is taken to be the largest likelihood of these removed $r$ samples and since $r$ new particles are sampled independently from the same distribution $\pi(\theta|l(\theta) > \epsilon_i)$, this is done in parallel. This results in a faster compression of the prior mass, the new particle $\theta^\star$ is sampled from the right distribution $\pi(\theta|l(\theta) > \epsilon_i)$ and the process is run in parallel. However, the authors in [6] argue that to achieve a similarly low variance of $t_r$ as for $t_1$ one needs to use a higher number of NS samples $N_r$ for the parallel NS algorithm with $r$ parallel processes, which compares to the number of NS samples $N_1$ used for sequential NS through $N_r \approx \sqrt{r}N_1$ particles.

#### S1.2 Numerical parallelization example

In section 2.2 we described our parallelization scheme. The difference to the parallelization schemes in [6] is how we weight the particles for the evidence approximation.

We denote with $t_j$ the random number that is distributed as the $j^{\text{th}}$ highest number among $N$ uniform numbers on the interval $[0, 1]$

Assume at the beginning of iteration $i$ the prior mass corresponding to the live particles is $x_{i-1,r}$. The prior volume shrinkage after removing $\theta_{i,1}$ is just the same as for regular nested sampling $x_{i,1} = t_1 x_{i-1,r}$, since $\theta_{i,1}$ is the particle with the lowest likelihood among $N$ uniformly distributed particles over $\pi(\theta|l(\theta) > \epsilon_{i-1,r})$. After removing the next particle $\theta_{i,2}$ the remaining the remaining prior volume is $x_{i,2} = t_2 x_{i-1,r}$. Thus, each prior volume can be written as $x_{i,j} = t_j x_{i-1,r}$ (with the obvious boundary condition $x_{0,r} = 1$). The variance of $t_r$ is monotonically increasing until $r = \frac{N+1}{2}$ and decreases then again, thus the variance of each $t_j$ can be upper bound by the variance of $t_r$ as long as $r \leq \frac{N+1}{2}$ (otherwise it can be upper bounded by the variance of $t_{r'}$ with $r' = \frac{N+1}{2}$).

We denote the Bayesian evidence approximation, using all samples $\epsilon_{i,j}$ up to $\epsilon_{m,k}$ with

$$\widetilde{Z}_{\mathcal{D}}^{m,k} = \sum_{i=1}^{m-1} \sum_{j=1}^{r} \epsilon_{i,j}(x_{i,j-1} - x_{i,j}) + \sum_{j=1}^{k} \epsilon_{m,j}(x_{m,j-1} - x_{m,j})$$

We illustrate the variance of the Bayesian evidence estimation on a small example. We assumed a likelihood function $l(\theta) = 100 \exp(-100\theta)$ with $\Omega = [0, 1]$, and ran the LF-NS algorithm for this example. In this example, the prior volume $x_{i,j}$ corresponding to the parameter $\theta_{i,j}$ is equal to this parameter $x_{i,j} = \theta_{i,j}$. In Figure S1 A we plotted the resulting values for $\widetilde{Z}_{\mathcal{D}}^{i,j}$ for each value $1 \leq i \leq m$ and $1 \leq j \leq r$. We ran $2000/r$

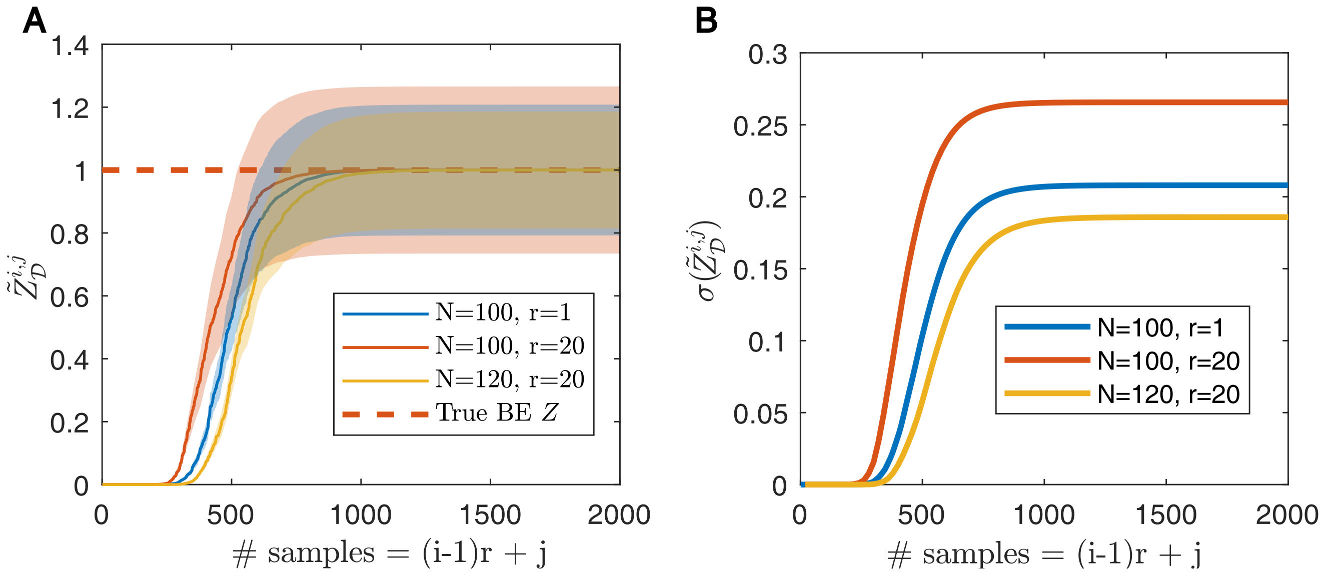

**Figure S1:** **A:** Values for $\widetilde{Z}_{\mathcal{D}}^{i,j}$ for LF-NS run. The shaded areas indicate the standard error. **B:** The standard deviation of $\widetilde{Z}$.

iterations for different values of $N$ and $r$. The resulting estimations of $\widetilde{Z}_{\mathcal{D}}^{i,j}$ and the corresponding standard deviations are shown in Figure S1 A and B. As can be seen when taking the same value for $N$, the parallel version with $r = 20$ has a slighter higher variance. However, when increasing the number of NS particles to $N_r = N + r = 120$ the variance decreases even compared to the sequential case ($r = 1$). Note that the speed up of the paralle version compared to the sequential version is a factor of $r$.

#### S2 Sampling from the super-level set

In the course of the LF-NS algorithm the parameter-likelihood pairs $\{\theta^\star, \widehat{l}^\star\} \in \Omega \times \mathbb{R}_{\geq 0}$ need to be sampled from distributions of the form

$$\Pi(\theta, \widehat{l}(\theta) | \widehat{l}(\theta) > \epsilon) = \pi(\theta) p(\widehat{l}(\theta) | \theta, \widehat{l}(\theta) > \epsilon).$$

Since the constrained distribution of the likelihood approximation $p(\widehat{l}(\theta)|\theta, \widehat{l}(\theta) > \epsilon)$ cannot be sampled directly, we resort to rejection sampling. The parameter vector $\theta^\star$ is sampled from $\pi(\theta)$ and the likelihood approximation $\widehat{l}^\star$ is sampled from $p(\widehat{l}(\theta)|\theta)$. The pair $\{\theta^\star, \widehat{l}^\star\}$ is accepted if $\widehat{l}^\star > \epsilon$ otherwise a new $\theta^\star$ and $\widehat{l}^\star$ are being sampled. The marginal distribution of $\theta$ is

$$\theta \sim \int \Pi(\theta, \widehat{l}(\theta) | \widehat{l}(\theta) > \epsilon_i) d\widehat{l}(\theta) = \pi(\theta) p(\widehat{l}(\theta) > \epsilon). \tag{2.1}$$

Since $p(\widehat{l}(\theta) > \epsilon)$ will in general be almost zero for large areas of $\Omega$, it is important to sample $\theta^\star$ not from the full prior $\pi(\theta)$ but rather just from the prior where $p(\widehat{l}(\theta) > \epsilon)$ is larger than zero. At each iteration $i$ of the LF-NS scheme the distribution 2.1 is already given through the $N - r$ live samples in the set $\mathcal{L}$, thus the challenge is to use these samples to sample uniformly from the prior on the support of $\pi(\theta)p(\widehat{l}(\theta) > \epsilon)$. This is similar to the sampling task in standard NS where instead of distribution 2.1, the set $\pi(\theta|l(\theta) > \epsilon)$ needs to be sampled. For NS the most popular ways to sample from these distributions are slice sampling [5] and ellipsoidal sampling [2]. Unfortunately, slice sampling cannot be applied to the case of LF-NS since unlike the likelihood function $l(\theta)$ in the NS case, the function $p(\widehat{l}(\theta))$ cannot be evaluated. Also, unlike the distribution $\pi(\theta|l(\theta) > \epsilon)$, which has sharp boarders, the distribution $\pi(\theta)p(\widehat{l}(\theta) > \epsilon)$ has smooth boundaries, making density estimation techniques more appropriate than ellipsoidal sampling. In the following we illustrate three ways of performing this sampling on a small example.

- **Ellipsoid sampler** This sampler was suggested in [8] and improved upon in [3] and is used in those paper to sample the distribution $\pi(\theta|l(\theta) > \epsilon)$. The basic idea is to create an ellipsoid that encloses all of the points in $\mathcal{L}$ and then sample from this ellipsoid uniformly. This ellipsoid is usually taken to be the one being spanned by the eigenvectors of the covariance matrix of the samples in $\mathcal{L}$, scaled in a way that it encompasses all points in $\mathcal{L}$. In [3] this approach was extended by first clustering the points in $\mathcal{L}$ and only then constructing ellipsoids for each cluster. Then the samples $\theta^\star$ were sampled uniformly from the union of these ellipsoids (in this case one has to take care not to oversample the intersections of these ellipsoids).

- **Kernel density estimation (KDE)** This approach estimates the density $p(\widehat{l}(\theta) > \epsilon)$ by placing a kernel $\mathcal{K}(\cdot|\theta_i)$ on each of the points $\theta_i$ in $\mathcal{L}$. To make sure that $\theta^\star$ is sampled uniformly from the support of $p(\widehat{l}(\theta) > \epsilon)$ each kernel is weighted by $w_i = \left(\frac{1}{|\mathcal{L}|}\sum_{j=1}^{|\mathcal{L}|}\mathcal{K}(\theta_i|\theta_j)\right)^{-1}$ and thus each new particle is sampled from

$$\theta^\star \sim \left(\sum_{j=1}^{|\mathcal{L}|} w_i\right)^{-1} \sum_{j=1}^{|\mathcal{L}|} w_i \mathcal{K}(\cdot|\theta_j). \tag{2.2}$$

  With this approach the samples $\theta^\star$ are not actually sampled uniformly from the support of $p(\widehat{l}(\theta) > \epsilon)$ but as the number of points in $\mathcal{L}$ the distribution of $\theta^\star$ approaches the uniform distribution.

- **Dirichlet process Gaussian mixture models (DP-GMM)** The is the approach we are following in this paper. DP-GMM approximates the distribution of the points in $\mathcal{L}$ through a mixture of Normal distributions $\widetilde{\mathcal{L}}$

$$\mathcal{L} \sim \widetilde{\mathcal{L}} = \sum_{j=1}^{k} w_j \mathcal{N}(\cdot|\mu_j, \Sigma_j),$$

  where the number $k$, mean $\mu_j$ and shape $\Sigma_j$ of each Gaussian are estimated from the data. This is done by placing a prior distribution on the Gaussian mixture shape and form and inferring the posterior of these parameters from the data. This inference is done through iterative Gibbs sampling. The details of the DP-GMM algorithm can be found in [4]. For our algorithm runs we implemented the algorithm from [4] in C++. To sample uniformly from the estimated density we employ rejection sampling where each $\theta^\star$ gets discarded right after sampling (before we even sample $\widehat{l^\star}$) with a probability of $(1 - \frac{g}{\widetilde{\mathcal{L}}(\theta^\star)})$ and $g$ is chosen to be the 0.1% quantile of $\widetilde{\mathcal{L}}$ (if $\widetilde{\mathcal{L}}(\theta^\star)$ is below the 0.1%, $\theta^\star$ gets also rejected). While this approach is computationally more demanding than the other two approaches, in our experience it provides by far the most reliable results. The computational overhead from the inference of $\widetilde{\mathcal{L}}$ and the rejection sampling of $\theta^\star$ was in all our examples negligible compared to the total computational effort.

In Figure S2 we illustrated the different mentioned sampling schemes. As an example we took the birth death example from the main paper but this time inferred both, $k$ and $\gamma$ (thus in this case we have $\theta = \{k, \gamma\}$). We approximated the likelihood $\widehat{l}(\theta)$ using $H = 20$ particle filter particles. Figure S2 A shows the true density $\pi(\theta)p(\widehat{l}(\theta) > \epsilon)$ for $\log(\epsilon) = -118.75$. This distribution was obtained by sampling $10^6$ particles from $\pi(\theta)p(\widehat{l}(\theta) > \epsilon)$ (which was done by sampling particles from $\pi(\theta)$ and accepting them if their approximated likelihood was above $\epsilon$). The red dots indicate the 90 particles in $\mathcal{L}$. For Figure S2 B we

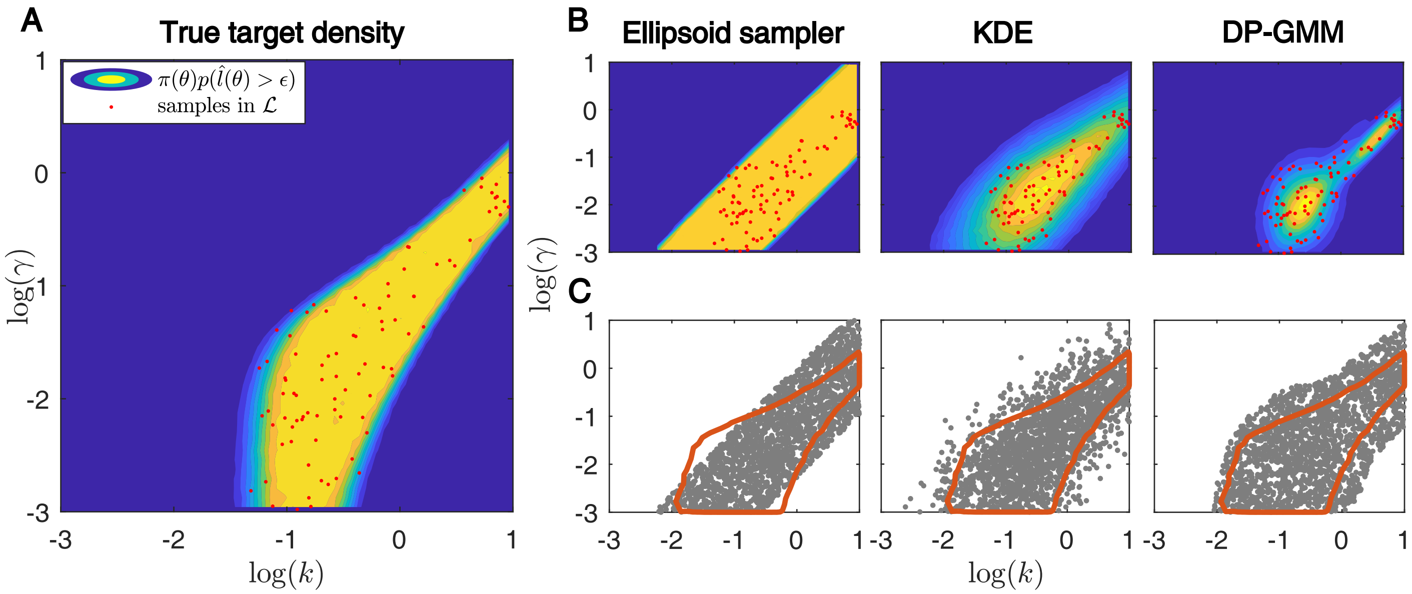

**Figure S2:** **A**: Contour lines of the distribution of $\pi(\theta)p(\widehat{l}(\theta) > \epsilon)$ for the birth death example, where $\theta = \{k, \gamma\}$, the number of particle filter particles to approximate $\widehat{l}(\theta)$ is $H = 20$ and $\log(\epsilon) = -118.75$. The density was approximated with $10^6$ samples. The red dots indicate 90 samples in $\mathcal{L}$. **B**: The estimations of $\mathcal{L}$ as obtained through an ellipsoid estimation, kernel density estimation (KDE) and Dirichlet process Gaussian mixture models (DP-GMM) based on the samples in $\mathcal{L}$. **C**: 2000 samples from the corresponding estimations of $\mathcal{L}$ where the samples for KDE were obtained according to 2.2 and the samples from DP-GMM were obtained using rejection sampling as described in section S2.

used the Ellipsoid sampler, KDE (where each Kernel was chosen to be a Gaussian with a covariance matrix equal to the empirical covariance matrix over all points in $\mathcal{L}$) and DP-GMM to approximate the set $\mathcal{L}$. As expected, the estimation with the ellipsoid encompasses all points in $\mathcal{L}$ but is in general not very tight. The KDE and DP-GMM estimations both provide a density estimation of the set $\mathcal{L}$. Figure S2 C finally shows how the obtained candidate particles $\theta^\star$ are distributed for each of the sampling methods. For the ellipsoid these particles are just uniformly sampled from the ellipsoid, while for KDE and DP-GMM the particles were obtained as described above to guarantee that they are uniformly distributed. The red line indicates the support of the target distribution $\pi(\theta)p(\widehat{l}(\theta) > \epsilon)$ (which was approximated by plotting the envelop of the $10^6$ samples from $\pi(\theta)p(\widehat{l}(\theta) > \epsilon)$).

We also tried to run the LF-NS method with the ellipsoid sampler and the KDE sampler on the LacGfp example, but in both cases we were not able to obtain a meaningful solution. With the ellipsoid sampler the acceptance rate dropped quickly very low ($\sim 10^{-5}$ and the computational time for each iteration became unreasonably long). While the LF-NS run with the KDE sampler converged, the approximated Bayesian evidence was three orders of magnitude below our estimate with DP-GMM and inspecting the marginal posterior distributions showed that the parameter space was clearly not explored fully.

##### S2.1 Final remarks regarding sampling for the contrained joint prior

We like to stress that each of the mentioned methods as well as other sampling methods such as MCMC samplers are suitable to sample from the constrained joint prior $\pi(\theta)p(\widehat{l}(\theta) > \epsilon)$. For each of these methods there exist plenty of variations that may prove particularly suitable for different cases. The brief outline in this section should be understood as a motivating illustration for the use of the DP-GMM sampler.

#### S3 Accuracy of the likelihood approximation $\widehat{l}$

As we have shown, LF-NS results in an unbiased estimation of the Bayesian evidence for any unbiased estimator of the likelihood $\widehat{l}(\theta)$. While the resulting Bayesian evidence does not depend on the particular

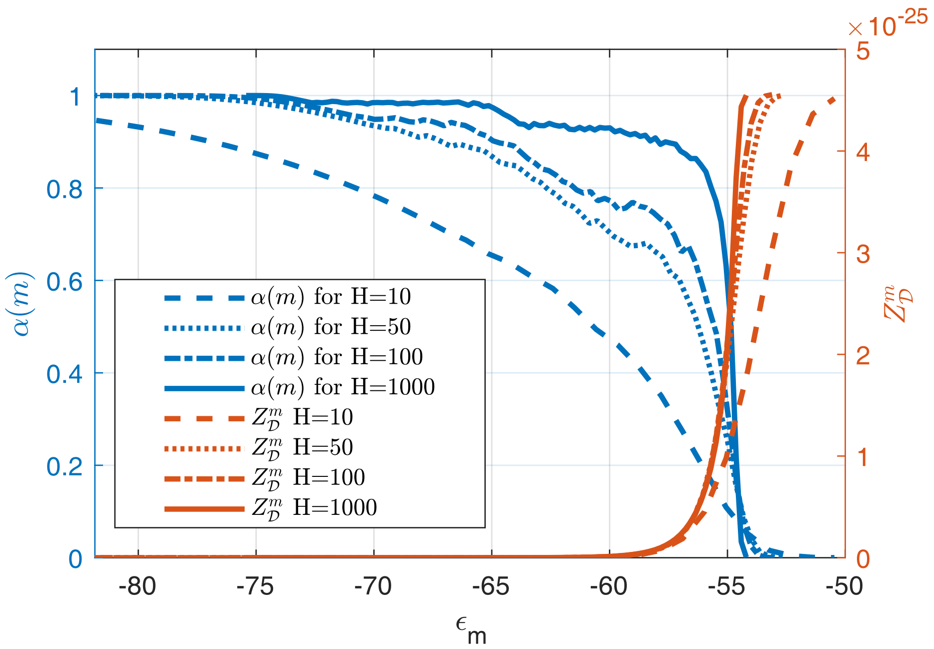

**Figure S3:** Acceptance rates $\alpha(m)$ and $Z_\mathcal{D}^m$ from $10^6$ samples of $\Pi(k, \hat{l}(k)$ for the birth death model for different values of particle filter particles $H$ and different iteration numbers $m$

likelihood approximation, the acceptance rate and thus the runtime of the LF-NS algorithm will in general depend on the variance of the likelihood approximation and the shape of the likelihood function. To illustrate this we proceed as in the main paper. We used the birth-death example and sampled $10^6$ particles from the distribution $\Pi(k, \hat{l}(k))$. For each of the $10^6$ samples $\{k_i, \hat{l}_i\}$ we set $\epsilon_i = l_i$ and considered the particles $k_i^- = \min(k_j : l_j \geq \epsilon_i)$ and $k^+ = \max(k_j : l_j \geq \epsilon_i)$. As in the main paper, wee denote with $S_i^+$ all the particles between $k_i^-$ and $k_i^+$ with a likelihood above $\epsilon_i$ and with $S_i^-$ the particles with a likelihood below $\epsilon_i$

$$S^+ = \{j : l_j > \epsilon_i, k_i^+ \geq k_j \geq k_i^-\} \quad \text{and} \quad S^- = \{j : l_j \leq \epsilon_i, k_i^+ \geq k_j \geq k_i^-\}$$

and computed the ratio of the number of their elements

$$\alpha(m) = \frac{|S_m^+|}{|S_m^-| + |S_m^+|}.$$

We repeated this for four different values for the number of particle filter particels $H = 10$, 50, 100 and 1000 and plotted the acceptance rates in Figure S3. The higher the number of particles $H$, the lower the resulting likelihood approximation variance. As expected, as $H$ increases, $\alpha(i)$ approaches the constant function 1. We also plotted the corresponding values for $Z_\mathcal{D}^m$ for different values of $H$ as an indicator of how much information about the final Bayesian evidence the particles in the prior volumes corresponding to each $\epsilon_m$ contain. The acceptance rate will vary from problem to problem and it is in general not possible to know ahead of time what level of accuracy of the likelihood approximation is necessary for what acceptance rate. However, a low acceptance rate is a strong indicator that the variance of the likelihood approximation needs to be decreased.

#### S4 Estimating the variance for the parallel LF-NS scheme

In the following we discuss in detail the approximation error for the Bayesian evidence

$$Z = \int_0^1 L(x)dx.$$

LF-NS approximates the above integral by approximating it on the final prior volume $x_{m,r}$ and the remaining volume separately

$$Z = \underbrace{\int_0^{x_{m,r}} L(x)dx}_{=:Z_{\mathcal{L}}^{m,r}} + \underbrace{\int_{x_{m,r}}^1 L(x)dx}_{=:Z_{\mathcal{D}}^{m,r}}.$$

The integral $Z_{\mathcal{D}}^{m,r}$ is approximated through a finite sum

$$\int_{x_{m,r}}^1 L(x)dx \approx \sum_{i=1}^m \sum_{j=1}^r L(x_{i,j})(x_{i,j-1} - x_{i,j}) = \sum_{i=1}^m \sum_{j=1}^r \epsilon_{i,j}(x_{i,j-1} - x_{i,j}) =: \widetilde{Z}_{\mathcal{D}}^{m,r}. \tag{4.3}$$

Since the prior volumes $x_{i,j}$ are in general not know, the numerical approximation $\widetilde{Z}_{\mathcal{D}}^{m,r}$ is approximated itself through the estimator

$$\widehat{Z}_{\mathcal{D}}^{m,r} = \sum_{i=1}^m \sum_{j=1}^r \epsilon_{i,j}(\widehat{x}_{i,j-1} - \widehat{x}_{i,j}) \approx \widetilde{Z}_{\mathcal{D}}^{m,r},$$

where the prior volumes $x_{i,j}$ are treated as random variables and approximated by their means $\widehat{x}_{i,j}$. The error in estimating $Z_{\mathcal{D}}$ through $\widetilde{Z}_{\mathcal{D}}^{m,r}$ is negligible compare to the error in estimating $\widetilde{Z}_{\mathcal{D}}^{m,r}$ through $\widehat{Z}_{\mathcal{D}}^{m,r}$ (see the discussion in [10] or [1]).

To emphasize its dependence on the final volume $x_{m,r}$, we rewrite the integral $Z_{\mathcal{L}}^{m,r}$ as an integral over the parameter space rather than the volume space

$$Z_{\mathcal{L}}^{m,r} = \int_0^{x_{m,r}} L(x)dx = x_{m,r} \underbrace{\int \widehat{l}(\theta) d\Pi(\theta, \widehat{l}(\theta) | \widehat{l}(\theta) \geq \epsilon_{mr})}_{=:L_{m,r}}$$

The quantity $L_{m,r}$ is the average of the likelihoods over the joint prior $\Pi(\theta, \widehat{l}(\theta))$ constraint to the likelihood regions above $\epsilon_{m,r}$. We approximate $L_{m,r}$ with a Monte Carlo estimator $\overline{L}_{m,r}$ and and the prior volume $x_{m,r}$ with its mean $\widehat{x}_{m,r}$

$$Z_{\mathcal{L}}^{m,r} \approx \widehat{x}_{m,r} \overline{L}_{m,r} =: \widehat{Z}_{\mathcal{L}}^{m,r}.$$

Thus, the final estimation error can be written as

$$\left\| Z - (\widehat{Z}_{\mathcal{D}}^{m,r} + \widehat{Z}_{\mathcal{L}}^{m,r}) \right\|$$

$$= \left\| x_{m,r} L_{m,r} - \widehat{Z}_{\mathcal{L}}^{m,r} + \int\limits_{x_{m,r}}^{1} L(x)dx - \widetilde{Z}_{\mathcal{D}}^{m,r} + \widetilde{Z}_{\mathcal{D}}^{m,r} - \widehat{Z}_{\mathcal{D}}^{m} \right\|$$

$$\leq \left\| \int\limits_{x_{m,r}}^{1} L(x)dx - \widetilde{Z}_{\mathcal{D}}^{m,r} \right\| + \left\| \underbrace{x_{m,r} L_{m,r} - \widehat{Z}_{\mathcal{L}}^{m,r}}_{=:\eta_{\mathcal{L}}^{m,r}} + \underbrace{\widetilde{Z}_{\mathcal{D}}^{m,r} - \widehat{Z}_{\mathcal{D}}^{m,r}}_{\eta_{\mathcal{D}}^{m,r}} \right\|.$$

The first part is the error from replacing the integral with a finite sum and, as mentioned before, is negligible. The errors $\eta_{\mathcal{L}}^{m,r}$ and $\eta_{\mathcal{D}}^{m,r}$ are all random variables where $\eta_{\mathcal{L}}^{m,r}$ represents the error of approximating $L_{m,r}$ with its Monte Carlo estimate $\overline{L}_{m,r}$ and replacing the final prior volume $x_{m,r}$ with its mean $\widehat{x}_{m,r}$ and the error $\eta_{\mathcal{D}}^{m,r}$ represents the error of estimating the random variables $x_{i,j}$ with their means $\widehat{x}_{i,j}$. Both errors have clearly mean 0.

As mentioned in the main paper, the prior volumes $x_{i,j}$ can be viewed as a product of the random variables $t_j$

$$x_{i,j} = t_j^{(i)} x_{i-1,r} = t_j^{(i)} \prod_{k=1}^{i-1} t_r^{(k)}.$$

The super script $(i)$ in $t_j^{(i)}$ emphasizes that it is the $i^{\text{th}}$ sample of the random variable $t_j$. The random variable $t_j$ is distributed as the $j^{\text{th}}$ highest number among $N$ uniform numbers on the interval $[0,1]$, which is the Beta distribution

$$t_j \sim \mathcal{B}(N-j+1, j)$$

with

$$\widehat{t}_j := \mathbb{E}(t_j) = \frac{N-j+1}{N+1}, \quad \mathbb{E}(t_j^2) = \frac{(N-j+1)(N-j+2)}{(N+1)(N+2)}$$

and variances of

$$\text{Var}\,(t_j) = \frac{(N-j+1)j}{(N+2)(N+1)^2}.$$

The corresponding values for the variables $x_{i,j}$ are

$$\widehat{x}_{i,j} := \mathbb{E}(x_{i,j}) = \widehat{t}_r^{i-1} \widehat{t}_j = \frac{(N-r+1)^{i-1}(N-j+1)}{(N+1)^i},$$

$$\mathbb{E}(x_{i,j}^2) = \mathbb{E}\left( t_j^{(i)^2} \prod_{k=1}^{i-1} t_r^{(k)^2} \right) = \mathbb{E}(t_j^2)\mathbb{E}(t_r^2)^{i-1} = \frac{(N-j+1)(N-j+2)(N-r+1)^{i-1}(N-r+2)^{i-1}}{(N+1)^i(N+2)^i}.$$

With this notation we compute the variances of the errors $\eta_L^m$, $\eta_{\mathcal{L}}^{m,r}$ and $\eta_{\mathcal{D}}^{m,r}$.

#### S4.1 Variance of $\eta_{\mathcal{L}}$

The error $\eta_{\mathcal{L}}$ is the error of approximating the final prior volume $x_{m,r}$ with its mean $\widehat{x}_{m,r}$ and replacing the integral $L_m$ with its Monte Carlo estimate $\overline{L}_m$ when approximating the Bayesian evidence over the final prior volume. Its variance is

$$\text{Var}(\eta_{\mathcal{L}}^{m,r}) = \text{Var}(x_{m,r} L_{m,r} - \widehat{x}_{m,r} \overline{L}_{m,r}) = \text{Var}\,(x_{m,r})\, L_{m,r}^2 + \text{Var}\left(\overline{L}_{m,r}\right) \widehat{x}_{m,r}^2.$$

The integral $L_{m,r}$ for the variance estimation can be estimated through the Monte Carlo estimate $\overline{L}_{m,r}$ and the variance of $\overline{L}_{m,r}$ is just

$$\operatorname{Var}\left(\overline{L}_{m,r}\right) = \frac{\sigma_L^2}{N}, \quad \sigma_L^2 = \frac{1}{N-1} \sum_{\{\theta,\widehat{l}\}\in\mathcal{L}_m} (\widehat{l} - \overline{L}_{m,r})^2.$$

The last average is taken over the points in the last live point set $\mathcal{L}_{m,r}$, which are all distributed according to $\Pi(\theta, \widehat{l}(\theta)|\widehat{l}(\theta) > \epsilon_{m,r})$.

#### S4.2 Variance of $\eta_{\mathcal{D}}$

Next, we estimate the variance of the estimation through the dead points. We have $\operatorname{Var}(\eta_{\mathcal{D}}) = \operatorname{Var}(\widetilde{Z}_{\mathcal{D}}^{m,r} - \widehat{Z}_{\mathcal{D}}^{m,r}) = \operatorname{Var}(\widetilde{Z}_{\mathcal{D}}^{m,r})$. We first rewrite

$$\widetilde{Z}_{\mathcal{D}}^{m,r} = \sum_{i=1}^{m}\sum_{j=1}^{r} \epsilon_{i,j}(x_{i,j-1} - x_{i,j}) = \sum_{i=1}^{m} x_{i-1,r} \underbrace{\sum_{j=1}^{r} \epsilon_{i,j}\left(t_{j-1}^{(i)} - t_j^{(i)}\right)}_{=:E_i}. \tag{4.4}$$

We point out that for each $i$, the samples $x_{i-1,r}$ and $E_i$ are statistically independent since $E_i$ only contain the $i^{\text{th}}$ samples of $t$ and $x_{i-1,r} = \prod_{k=1}^{i-1} t_r^{(k)}$ contains only the samples up to $i-1$. For $m > 1$[1] we can write

$$\operatorname{Var}(\widetilde{Z}_{\mathcal{D}}^{m,r}) = \operatorname{Var}(\widetilde{Z}_{\mathcal{D}}^{m-1,r}) + 2\operatorname{Cov}(x_{m-1,r}E_m, \widetilde{Z}_{\mathcal{D}}^{m-1,r}) + \operatorname{Var}(x_{m-1,r}E_m). \tag{4.5}$$

We fist compute the general formula for $\operatorname{Cov}\left(x_m, \widetilde{Z}_{\mathcal{D}}^{m,r}\right)$. The product of $x_{m,r}$ and $\widetilde{Z}_{\mathcal{D}}^{m,r}$ is

$$x_{m,r}\widetilde{Z}_{\mathcal{D}}^{m,r} = \sum_{i=1}^{m} x_{m,r}x_{i-1,r}E_i$$

$$= \sum_{i=1}^{m}\prod_{l=1}^{i-1} t_r^{(l)^2} \prod_{k=i+1}^{m} t_r^{(k)} \sum_{j=1}^{r} \epsilon_{i,j}\left(t_r^{(i)}t_{j-1}^{(i)} - t_r^{(r)}t_j^{(i)}\right).$$

In the above formulation we used the convention that $\prod_{k=m}^{m-1} = 1$. With this and using that for random variables $X$, $Y$ and $Z$ we have $\mathbb{E}\left(X^2\right)\mathbb{E}\left(YZ\right) - \mathbb{E}\left(X\right)^2\mathbb{E}\left(Y\right)\mathbb{E}\left(Z\right) = \operatorname{Var}\left(X\right)\mathbb{E}(YZ) + \mathbb{E}(X)^2\operatorname{Cov}\left(Y,Z\right)$ we compute the covariance

$$\operatorname{Cov}(x_m, \widetilde{Z}_{\mathcal{D}}^{m}) = \sum_{i=1}^{m} \widehat{x}_{m-i,r} \sum_{j=1}^{r} \epsilon_{i,j}\left(\operatorname{Var}(x_{i-1,r})(\mathbb{E}(t_r t_{j-1}) - \mathbb{E}(t_r t_j)) + \widehat{x}_{i-1,r}^2(\operatorname{Cov}(t_r, t_{j-1}) - \operatorname{Cov}(t_r, t_j))\right) \tag{4.6}$$

[1]for $m = 1$ we obviously have $\operatorname{Var}\left(\widetilde{Z}_{\mathcal{D}}^1\right) = \operatorname{Var}\left(E_1\right)$.

$$= \sum_{i=1}^{m} \widehat{x}_{m-i,r} \sum_{j=1}^{r-1} \epsilon_{i,j} \operatorname{Var}\left(x_{i-1,r}\right) \left(\widehat{t}_r \widehat{t}_{j-1} - \widehat{t}_r \widehat{t}_j\right))$$

$$+ \sum_{i=1}^{m} \widehat{x}_{m-i,r} \epsilon_{i,r} \left(\operatorname{Var}\left(x_{i-1,r}\right) \widehat{t}_r \widehat{t}_{r-1} - \operatorname{Var}\left(x_{i,r}\right)\right).$$

Next we compute the variance

$$\operatorname{Var}(x_{m-1,r} E_m) = \mathbb{E}\left(x_{m-1}^2\right) \operatorname{Var}(E_m) + \operatorname{Var}(x_{m-1,r}) \widehat{E}_m^2. \tag{4.7}$$

The square of $E_m$ is

$$E_m^2 = \left(\sum_{j=1}^{r} \epsilon_{m,j} \left(t_{j-1}^{(m)} - t_j^{(m)}\right)\right)^2$$

$$= \sum_{j=1}^{r} \sum_{k=1}^{r} \epsilon_{m,j} \epsilon_{m,k} \left(t_{j-1}^{(m)} - t_j^{(m)}\right) \left(t_{k-1}^{(m)} - t_k^{(m)}\right)$$

and thus the variance of $E_m$ is

$$\operatorname{Var}(E_m) = \sum_{j=1}^{r} \sum_{k=1}^{r} \epsilon_{m,j} \epsilon_{m,k} \operatorname{Cov}\left(t_{j-1} - t_j, t_{k-1} - t_k\right).$$

Putting this variance of $E_m$ into formula 4.7 we obtain

$$\operatorname{Var}(x_{m-1,r} E_m) =$$

$$= \sum_{j=1}^{r} \sum_{k=1}^{r} \epsilon_{m,j} \epsilon_{m,k} \left(\mathbb{E}(x_{m-1,r}^2) \operatorname{Cov}\left(t_{j-1} - t_j, t_{k-1} - t_k\right) + \operatorname{Var}(x_{m-1,r}) \left(\widehat{t}_{j-1} - \widehat{t}_j\right) \left(\widehat{t}_{k-1} - \widehat{t}_k\right)\right)$$

$$= \sum_{j=1}^{r} \epsilon_{m,j}^2 \operatorname{Var}\left(x_{m-1,j-1} - x_{m-1,j}\right) + 2 \sum_{j=2}^{r} \sum_{k=1}^{j-1} \epsilon_{m,j} \epsilon_{m,k} (\operatorname{Var}\left(x_{m-1,r}\right) \left(1 - t_1\right))$$

$$-2 \sum_{j=2}^{r} \epsilon_{m,j} \epsilon_{m,j-1} \mathbb{E}\left(x_{m-1,r}^2\right) \operatorname{Var}\left(t_{j-1}\right)$$

Combing the variance and covariance term in 4.5 we get

$$\operatorname{Var}(\eta_{\mathcal{D}}^{m,r}) = \operatorname{Var}(\eta_{\mathcal{D}}^{m-1}) + 2\widehat{E}_m \operatorname{Cov}(x_{m-1}, \widetilde{Z}_{\mathcal{D}}^{m-1}) + \operatorname{Var}(x_{m-1,r} E_m) \tag{4.8}$$

##### S4.3 Total variance

The total variance is

$$\mathrm{Var}(\eta_{\mathcal{L}}^{m,r} + \eta_{\mathcal{D}}^{m,r}) = \mathrm{Var}\left(\eta_{\mathrm{L}}^{m,r}\right) + \mathrm{Var}\left(\eta_{\mathcal{D}}^{m,r}\right) + 2\,\mathrm{Cov}\left(\eta_{\mathcal{L}}^{m,r}, \eta_{\mathcal{D}}^{m,r}\right).$$

Since $\eta_{\mathcal{L}}$ and $\eta_{\mathcal{D}}$ are not independent (they both share the random variable $x_{m,r}$), we have to account for their dependence.

$$\mathrm{Cov}(\eta_{\mathcal{L}}^{m,r}, \eta_{\mathcal{D}}^{m,r}) = \mathrm{Cov}\left(x_{m,r} L_{m,r}, \widetilde{Z}_{\mathcal{D}}^{m,r}\right) = L_{m,r}\,\mathrm{Cov}\left(x_{m,r}, \widetilde{Z}_{\mathcal{D}}^{m,r}\right)$$

This covariance can be computed using formula 4.6. The total variance is

$$\mathrm{Var}(\eta_{\mathcal{L}}^{m,r} + \eta_{\mathcal{D}}^{m,r})$$

$$= \mathrm{Var}(\eta_{\mathcal{L}}^{m,r}) + \mathrm{Var}(\eta_{\mathcal{D}}^{m,r}) + 2L_{m,r}\,\mathrm{Cov}\left(x_{m,r}, \widetilde{Z}_{\mathcal{D}}^{m,r}\right)$$

#### S5 Lower bound on the variance $\mathrm{Var}(\eta_{\mathcal{L}}^{m,r} + \eta_{\mathcal{D}}^{m,r})$

The goal of this section is to establish for each iteration $m$ a lower bound for the total variance of the NS estimator. We start by introducing some notation for the variation of the error estimate. The double index $(m, k)$ indicates the corresponding quantities using all samples up until the $(m, k)$ so for instance

$$\widetilde{Z}_{\mathcal{D}}^{(m,k)} = \sum_{i=1}^{m-1} \sum_{i=1}^{r} \epsilon_{i,j}(x_{i,j-1} - x_{i,j}) + \sum_{i=1}^{k} \epsilon_{m,j}(x_{m,j-1} - x_{m,j}).$$

We continue to write

$$\sigma_{\mathrm{tot}}^{2m,j} := \mathrm{Var}(\eta_{\mathcal{L}}^{m,j} + \eta_{\mathcal{D}}^{m,j}).$$

Writing this formula out we get

$$\sigma_{\mathrm{tot}}^{2m,j} := \mathrm{Var}\left(x_{m,j}\right) L_{m,j}^2 + \mathrm{Var}\left(\overline{L}_{m,j}\right) \widehat{x}_{m,j}^2 + \mathrm{Var}\left(\eta_{\mathcal{D}}^{m,j}\right) + 2L_{m,j}\,\mathrm{Cov}\left(x_{m,r}, \widetilde{Z}_{\mathcal{D}}^{m,j}\right).$$

We define with $\sigma_{\mathrm{min}}^{2m}$ the same variance, but where we assume $\mathrm{Var}\left(\overline{L}_{m,j}\right) = 0$

$$\sigma_{\mathrm{min}}^{2m,j} := \mathrm{Var}\left(x_{m,j}\right) L_{m,j}^2 + \mathrm{Var}\left(\eta_{\mathcal{D}}^{m,j}\right) + 2L_{m,j}\,\mathrm{Cov}\left(x_{m,r}, \widetilde{Z}_{\mathcal{D}}^{m,j}\right).$$

Clearly, $\sigma_{\mathrm{tot}}^{2m,j}$ is monotonically increasing in $\mathrm{Var}\left(\overline{L}_{m,j}\right)$ and we have for each $m$ and each $j$

$$\sigma_{\mathrm{tot}}^{2m,j} \geq \sigma_{\mathrm{min}}^{2m,j}.$$

This is not surprising since this statement only says that the total variance at each iteration monotonically increases with the variance of the Monte Carlo estimator. As a second step we show that for all $m' \geq m$ we have

$$\sigma_{\mathrm{min}}^{2m',j} \geq \sigma_{\mathrm{min}}^{2m,j} \tag{5.9}$$

and for all $r \geq j' \geq j$ we have

$$\sigma_{\text{min}}^{m,j'} \geq \sigma_{\text{min}}^{m,j} \tag{5.10}$$

This statement means that assuming we know the integral of the likelihood function over the final prior volume (and thus the variance of the estimator $\overline{L}_m$ is 0), the variance of the LF-NS estimate only increases with further LF-NS iterations. Once we show this, we have shown that $\sigma_{\text{min}}^{2m,r}$ is a lower bound on the variance $\sigma_{\text{tot}}^{2m',r}$ for each iteration $m' \geq m$.

#### S5.1 Recursive formulations for variance terms

We show that $\sigma_{\text{min}}^{2m,j}$ is monotonically increasing in $j$. In the following it is understood that the double index $(m, 0) = (m-1, r)$. The variance $\sigma_{\text{min}}^{2m,j}$ is

$$\sigma_{\text{min}}^{2m,j} = \operatorname{Var}(x_{m,j} L_{m,j}) + \operatorname{Var}\left(\eta_{\mathcal{D}}^{m,j}\right) + 2\operatorname{Cov}\left(x_{m,r} L_{m,j}, \widetilde{Z}_{\mathcal{D}}^{m,j}\right)$$

We first write down the recursive formulas for all three involved terms.

- $\operatorname{Var}\left(\eta_{\mathcal{D}}^{m,j}\right)$:

$$\operatorname{Var}\left(\widehat{Z}_{\mathcal{D}}^{m,j}\right) = \operatorname{Var}\left(\widetilde{Z}_{\mathcal{D}}^{m,j-1} + \epsilon_{m,j}(x_{m,j-1} - x_{m,j})\right)$$

$$= \operatorname{Var}\left(\eta_{\mathcal{D}}^{m,j-1}\right) + \epsilon_{m,j}^2 \operatorname{Var}(x_{m,j-1} - x_{m,j}) + 2\epsilon_{m,j} \operatorname{Cov}\left(x_{m,j-1} - x_{m,j}, \widetilde{Z}_{\mathcal{D}}^{m,j-1}\right).$$

- $\operatorname{Var}(x_{m,j} L_{m,j})$: We write

$$x_{m,j} L_{m,j} = x_{m,j-1} L_{m,j-1} - (x_{m,j-1} - x_{m,j}) l_{m,j},$$

  where $l_{m,j}$ is the average of the likelihood over the areas where the likelihood is between $\epsilon_{m-1,j}$ and $\epsilon_{m,j}$

$$l_{m,j} = \int \widehat{l}(\theta) d\Pi(\theta, \widehat{l}(\theta) | \epsilon_{m,j} \geq \widehat{l}(\theta) \geq \epsilon_{m,j-1}).$$

  This in particular implies $\epsilon_{m,j} \geq l_{m,j} \geq \epsilon_{m,j-1}$. We obtain the recursive formula

$$\operatorname{Var}(x_{m,j} L_{m,j}) = \operatorname{Var}(x_{m,j-1} L_{m,j-1} - (x_{m,j-1} - x_{m,j}) l_{m,j})$$

$$= \operatorname{Var}(x_{m,j-1} L_{m,j-1}) + \operatorname{Var}((x_{m,j-1} - x_{m,j}) l_{m,j}) - 2 L_{m,j-1} l_{m,j} \operatorname{Cov}(x_{m,j-1} - x_{m,j}, x_{m,j-1})$$

- $\operatorname{Cov}\left(x_{m,j} L_{m,j}, \eta_{\mathcal{D}}^{m,j}\right)$:

$$\operatorname{Cov}\left(x_{m,j} L_{m,j}, \eta_{\mathcal{D}}^{m,j}\right) = \operatorname{Cov}\left(x_{m,j} L_{m,j}, \widetilde{Z}_{\mathcal{D}}^{m,j}\right)$$

$$= \operatorname{Cov}\left(x_{m,j-1} L_{m,j-1}, \eta_{\mathcal{D}}^{m-1}\right) + \operatorname{Cov}(x_{m,j-1} L_{m,j-1}, \epsilon_{m,j}(x_{m,j-1} - x_{m,j}))$$

$$- \operatorname{Cov}\left((x_{m,j-1} - x_{m,j}) l_m, \widetilde{Z}_{\mathcal{D}}^{m,j-1}\right) - \operatorname{Cov}((x_{m,j-1} - x_{m,j}) l_m, \epsilon_{m,j}(x_{m,j-1} - x_{m,j})).$$

$$= \mathrm{Cov}\left(x_{m,j-1}L_{m,j-1}, \eta_{\mathcal{D}}^{m,j-1}\right)$$

$$+L_{m,j-1}\epsilon_{m,j}\,\mathrm{Cov}\left(x_{m,j-1}-x_{m,j}, x_{m,j-1}\right) - l_{m,j}\,\mathrm{Cov}\left(x_{m,j-1}-x_{m,j}, \widetilde{Z}_{\mathcal{D}}^{m,j-1}\right)$$

$$-\epsilon_{m,j}l_m\,\mathrm{Var}\left(x_{m,j-1}-x_{m,j}\right).$$

#### S5.2 Lower bounding the residual

To obtain the full residue $\sigma_{\min}^{2m,j} - \sigma_{\min}^{2m,j-1}$ we combine the above computed residual terms:

$$\sigma_{\min}^{2m,j} - \sigma_{\min}^{2m,j-1}$$

$$= \epsilon_{m,j}^2\,\mathrm{Var}\left(x_{m,j-1}-x_{m,j}\right) + 2\epsilon_{m,j}\,\mathrm{Cov}\left(x_{m,j-1}-x_{m,j}, \widetilde{Z}_{\mathcal{D}}^{m,j-1}\right)$$

$$+l_{m,j}^2\,\mathrm{Var}\left(x_{m,j-1}-x_{m,j}\right) - 2L_{m,j-1}l_{m,j}\,\mathrm{Cov}\left(x_{m,j-1}-x_{m,j}, x_{m,j-1}\right)$$

$$+2L_{m,j-1}\epsilon_{m,j}\,\mathrm{Cov}\left(x_{m,j-1}-x_{m,j}, x_{m,j-1}\right) - 2l_{m,j}\,\mathrm{Cov}\left(x_{m,j-1}-x_{m,j}, \widetilde{Z}_{\mathcal{D}}^{m,j-1}\right)$$

$$-2\epsilon_{m,j}l_m\,\mathrm{Var}\left(x_{m,j-1}-x_{m,j}\right)$$

We group the terms

$$2(\epsilon_{m,j}-l_{m,j})\,\mathrm{Cov}\left(x_{m,j-1}-x_{m,j}, \widetilde{Z}_{\mathcal{D}}^{m,j-1}\right)$$

$$+2(\epsilon_{m,j}-l_{m,j})L_{m,j-1}\,\mathrm{Cov}\left(x_{m,j-1}-x_{m,j}, x_{m,j-1}\right)$$

$$+(\epsilon_{m,j}-l_{m,j})^2\,\mathrm{Var}\left(x_{m,j-1}-x_{m,j}\right)$$

Since we are only interested in showing that the residual is larger than zero, it is enough to show that the sum of the first two terms is larger than zero. For ease of computation we note that for $i \leq m$ and $j < r$

$$\mathrm{Cov}\left(x_{m,j-1}-x_{m,j}, x_{i-1,r}\right) = \mathrm{Cov}\left(x_{i-1,r}\prod_{k=i}^{m-1} t_r^{(k)}(t_{j-1}^{(m)} - t_j^{(m)}), x_{i-1,r}\right) = \widehat{x}_{m-i,r}\frac{1}{N+1}\,\mathrm{Var}\left(x_{i-1,r}\right)$$

and write for $j \leq r$

$$E_i^{(j)} = \sum_{k=1}^{j}\epsilon_{i,k}\left(t_{k-1}^{(i)} - t_k^{(i)}\right)$$

and similarly $\widehat{E}_i^{(j)}$ when we replace the $t_{k-1}^{(i)}$ and $t_k^{(i)}$with their means $\widehat{t}_{k-1}$ and $\widehat{t}_k$ respectively. We exploit the fact that $\epsilon_{m,j} \geq l_{m,j}$ and ignore the factor $2(\epsilon_{m,j}-l_{m,j})$. We proceed to write

$$\mathrm{Cov}\left(x_{m,j-1}-x_{m,j}, \widetilde{Z}_{\mathcal{D}}^{m,j-1}\right) + L_{m,j-1}\,\mathrm{Cov}\left(x_{m,j-1}-x_{m,j}, x_{m,j-1}\right)$$

$$= \sum_{i=1}^{m-1}\mathrm{Cov}\left(x_{m,j-1}-x_{m,j}, x_{i-1,r}E_i^{r-1}\right) + \mathrm{Cov}\left(x_{m,j-1}-x_{m,j}, x_{i-1,r}\epsilon_{i,r}t_{r-1}^{(i)}\right) - \mathrm{Cov}\left(x_{m,j-1}-x_{m,j}, x_{i-1,r}\epsilon_{i,r}t_r^{(i)}\right)$$

$$+\operatorname{Cov}\left(x_{m,j-1}-x_{m,j},x_{m-1,r}E_m^{j-2}\right)+\operatorname{Cov}\left(x_{m,j-1}-x_{m,j},x_{m-1,r}\epsilon_{m,j-1}t_{j-2}^{(m)}\right)-\operatorname{Cov}\left(x_{m,j-1}-x_{m,j},x_{m,j-1}\epsilon_{m,j-1}\right)$$

$$+L_{m,j-1}\operatorname{Cov}\left(x_{m,j-1}-x_{m,j},x_{m,j-1}\right)$$

$$=\sum_{i=1}^{m-1}\widehat{x}_{m-i,r}\frac{1}{N+1}\operatorname{Var}\left(x_{i-1,r}\right)\left(\widehat{E}_i^{r-1}+\epsilon_{i,r}\widehat{t}_{r-1}\right)-\widehat{x}_{m-i-1,r}\frac{1}{N+1}\operatorname{Var}\left(x_{i,r}\right)\epsilon_{i,r}$$

$$+\frac{1}{N+1}\operatorname{Var}\left(x_{m-1,r}\right)\left(\widehat{E}_m^{j-2}+\epsilon_{m,j-1}\widehat{t}_{j-2}\right)$$

$$+\underbrace{\left(L_{m,j-1}-\epsilon_{m,j-1}\right)\operatorname{Cov}\left(x_{m,j-1}-x_{m,j},x_{m,j-1}\right)}_{\geq 0}$$

We use that $\operatorname{Var}\left(x_{0,r}\right)=0$ and $\widehat{E}_i^{(j)}+\epsilon_{i,j+1}\widehat{t}_j\geq\epsilon_{i,1}$ and write

$$\sigma_{\min}^{2m,j}-\sigma_{\min}^{2m,j-1}\geq\sum_{i=2}^{m}\widehat{x}_{m-i,r}\frac{1}{N+1}\operatorname{Var}\left(x_{i-1,r}\right)\epsilon_{i,1}-\widehat{x}_{m-i,r}\frac{1}{N+1}\operatorname{Var}\left(x_{i-1,r}\right)\epsilon_{i-1,r}\geq 0$$

With this we have shown that $\sigma_{\min}^{2m,j+1}\geq\sigma_{\min}^{2m,j}$, which means that assuming that at iteration $m$ the integral $L_{m,r}$ over the final prior volume is known, each additional NS iteration only increases the variance. Thus $\sigma_{\min}^{2m,r}$ is a lower bound for the minimal achievable variance (using the same number $N$ of LF-NS particles).

### S6 Examples used

#### S6.1 Birth Death Model

As described in the main paper, the birth death model consists of a single species (mRNA) that gets created at a rate $k$ and degraded at a rate $\gamma$. This gives us two reactions

1. $\emptyset\xrightarrow{k}\text{mRNA}$ Transcription of mRNA.
2. $\text{mRNA}\xrightarrow{\gamma}\emptyset$ Degradation of mRNA.

We assume measurements $y_\tau=\mathcal{N}(\text{mRNA}(t_\tau),\sigma)$ with $\sigma=2$. For the example in the paper we fixed the value for $\gamma=1$.

#### S6.2 Lac-Gfp example

The second model we use for the demonstration of our algorithm is the rather large Lac-Gfp system with 9 species and 18 reactions. Table S1 shows the species involved and their initial distribution for the simulation.

**Table S1:** Species and intial numbers of the Lac-Gfp model

| Species | Notation | Initial Distribution |
|---|---|---|
| LacI mRNA | $lacI$ | U([0, 5]) |
| LacI protein monomer | $LACI$ | U([0, 10]) |
| LacI dimer | $LACI2$ | fixed to 0 |
| Unoccupied (active) Lac promoter | $PLac$ | fixed to 0 |
| Occupied Lac promoter with 2 repressor molecules bound | $O2Lac$ | fixed to 0 |
| Occupied Lac promoter with 4 repressor molecules bound | $O4Lac$ | U([50, 70]) |
| GFP mRNA | $gfp$ | fixed to 0 |
| "Dark" GFP protein | $GFP$ | fixed to 0 |
| Mature GFP protein | $mGFP$ | fixed to 0 |

U([$a, b$]) denotes the uniform distribution on the interval [$a, b$].

The reactions of the model all follow mass action kinetics and take the following form:

1. $\emptyset \xrightarrow{\theta_1} lacI$ Transcription of lacI mRNA (constitutive).
2. $lacI \xrightarrow{\theta_2} \emptyset$ Degradation of lacI mRNA (constitutive).
3. $lacI \xrightarrow{\theta_3} lacI + LACI$ Translation of LACI protein.
4. $LACI \xrightarrow{\theta_u} \emptyset$ where $\theta_u = \theta_4 + \theta_5[IPTG]$. Degradation of LACI protein, increased by the input (IPTG).
5. $LACI + LACI \xrightarrow{\theta_6} LACI2$ Dimerization of LACI protein.
6. $LACI2 \xrightarrow{\theta_7} LACI + LACI$ Dissociation of LACI dimer.
7. $LACI2 + PLac \xrightarrow{\theta_8} O2Lac$ Binding of LACI dimer to Lac operator sequence.
8. $O2Lac \xrightarrow{\theta_9} LACI2 + PLac$ Dissociation of LACI dimer from operator sequence.
9. $O2Lac + O2Lac \xrightarrow{\theta_{10}} O4Lac$ Binding of two LacI/operator complexes and tetramerization.
10. $O4Lac \xrightarrow{\theta_{11}} O2Lac + O2Lac$ Dissociation of tetramer structure.
11. $PLac \xrightarrow{\theta_{12}} PLac + gfp$ . Transcription of gfp mRNA from active Lac promoter.
12. $O2Lac \xrightarrow{\theta_{13}} O2Lac + gfp$ Transcription of gfp mRNA from Lac promoter bound to LacI dimer.
13. $O4Lac \xrightarrow{\theta_{14}} O4Lac + gfp$ Transcription of gfp mRNA from Lac promoter bound to LacI tetramer.
14. $gfp \xrightarrow{\theta_{15}} \emptyset$ Degradation of gfp mRNA.
15. $gfp \xrightarrow{\theta_{16}} gfp + GFP$ Translation of dark GFP protein.
16. $GFP \xrightarrow{\theta_{17}} \emptyset$ Degradation of dark GFP protein.
17. $GFP \xrightarrow{\theta_{18}} mGFP$ Maturation of GFP.
18. $mGFP \xrightarrow{\theta_{17}} \emptyset$ Degradation of mature GFP protein.

The effect of IPTG on the system is modelled as an increase in the degradation rate of $LACI$. The relationship between such rate and the inducer concentration, denoted $[IPTG]$, is assumed to be linear. We take the $[IPTG]$ concentration to be 10 $\mu M$.
The parameters used to simulate the dataset $\boldsymbol{y}$ as well as the priors used for the inference are shown in Table S2. We assume that our observation $y_\tau$ at a certain time $\tau$ is distributed according to

$$y_\tau \sim \mathcal{N}(22x_\tau, 5\sqrt{x_\tau}) + \mathcal{B},$$

where $x_\tau$ is the number of GFP molecules at time $t_\tau$, $\mathcal{N}(\mu, \sigma)$ is the normal distribution with mean $\mu$ and standard deviation $\sigma$ and $\mathcal{B}$ is a known background fluorescence assumed to be $\mathcal{B} = \mathcal{N}(80, 40)$. This noise model implies a mean fluorescence of 22 for each protein and a standard deviation of 5. These model choices for the simulation are inspired by the inferred parameters for the real biological data from [7].

**Table S2:** Prior distributions of the Lac-GFP parameters and the real values $\boldsymbol{\theta}^*$ used for the simulation of $\boldsymbol{y}$.

| Parameter | Meaning | Prior interval | $\boldsymbol{\theta}^*$ |
|---|---|---|---|
| $\theta_1$ | lacI transcription rate | $[10^{-5}, 10]$ | 1.5 |
| $\theta_2$ | lacI degradation rate | $[10^{-5}, 10]$ | 7.5 |
| $\theta_3$ | LACI translation rate | $[10^{-5}, 10]$ | 1.5 |
| $\theta_4$ | IPTG-independent LACI degradation rate | $[10^{-5}, 10]$ | 4.5 |
| $\theta_5$ | IPTG-induced increase in LACI degradation rate | $[10^{-5}, 10]$ | 5 |
| $\theta_6$ | Dimerization rate of LACI | [0.1, 3000] | 1650 |
| $\theta_7$ | Dissociation rate of LACI dimers | $[10^{-5}, 10]$ | 6 |
| $\theta_8$ | Binding rate of LACI dimers to Lac promoter | $[10^{-5}, 10]$ | 0.48 |
| $\theta_9$ | Dissociation rate of LACI dimers from Lac promoter | $[10^{-5}, 1]$ | 0.5 |
| $\theta_{10}$ | Tetramerization rate of LACI | [0.01, 500] | 230 |
| $\theta_{11}$ | Dissociation rate of LACI tetramers | $[10^{-5}, 10]$ | 0.4 |
| $\theta_{12}$ | gfp transcription rate from free PLac promoter | [0.01, 500] | 125 |
| $\theta_{13}$ | gfp transcription rate from LACI dimer-bound PLac | $[10^{-5}, 10]$ | 0.2 |
| $\theta_{14}$ | gfp transcription rate from LACI tetramer-bound PLac | $[10^{-5}, 1]$ | 0.01 |
| $\theta_{15}$ | gfp degradation rate | $[10^{-5}, 10]$ | 1.5 |
| $\theta_{16}$ | GFP translation rate | [0.01, 50] | 32 |
| $\theta_{17}$ | GFP degradation rate | $[10^{-5}, 10]$ | 1 |
| $\theta_{18}$ | GFP maturation rate | $[10^{-5}, 10]$ | 2.2 |

For all the Lac-Gfp inference problems presented in this paper, each parameter was assigned an independent uniform log prior distribution in the interval listed in the table.

##### S6.2.1 Likelihood approximation for the Lac-Gfp system

Figure S4 A shows the first 3 simulated trajectories of the Lac-Gfp system measured on 29 timepoints. As can be seen, most trajectories exhibit switch like behaviour. This is indeed one of the properties of the considered example that make it particularly hard to perform likelihood approximation for the single trajectories. Figure S4 B shows the distribution of 100 runs of particle filter for the estimation of the log-likelihood using different numbers $H$ of particles for the particle filter. As can be seen, the likelihood approximations vary over many thoursands of orders of magnitude even with as many as $H = 5000$ particles. Observe that while the likelihood approximation is unbiased, this will usually not hold for the log-likelihood approximation.

#### S6.3 Transcriptional model

The third example was taken from [9] and consists of a random promoter binding at a rate $k_{\text{on}}$ to a gene and turning it "on" and randomly unbinding from that same gene turning it "off" at a rate $k_{\text{off}}$. While in the "on" state, mRNA gets transcribed from that gene and can be measured as it is being transcribed. Due to the length of the gene the authors assume that the transcription takes about 2 minutes. Our model thus consists of the two species "$g_{\text{on}}$" and ''$g_{\text{off}}$" that switch between each other with the rates $k_{\text{on}}$ and $k_{\text{off}}$. To account for the fact that each nascant RNA gets observed as long as it is being transcribed, we introduce 8 virtual RNA species that transform from one to another at rate $\lambda$. As initial state, we pick $g_{\text{off}} = 1$ and all other species as 0. The reactions are

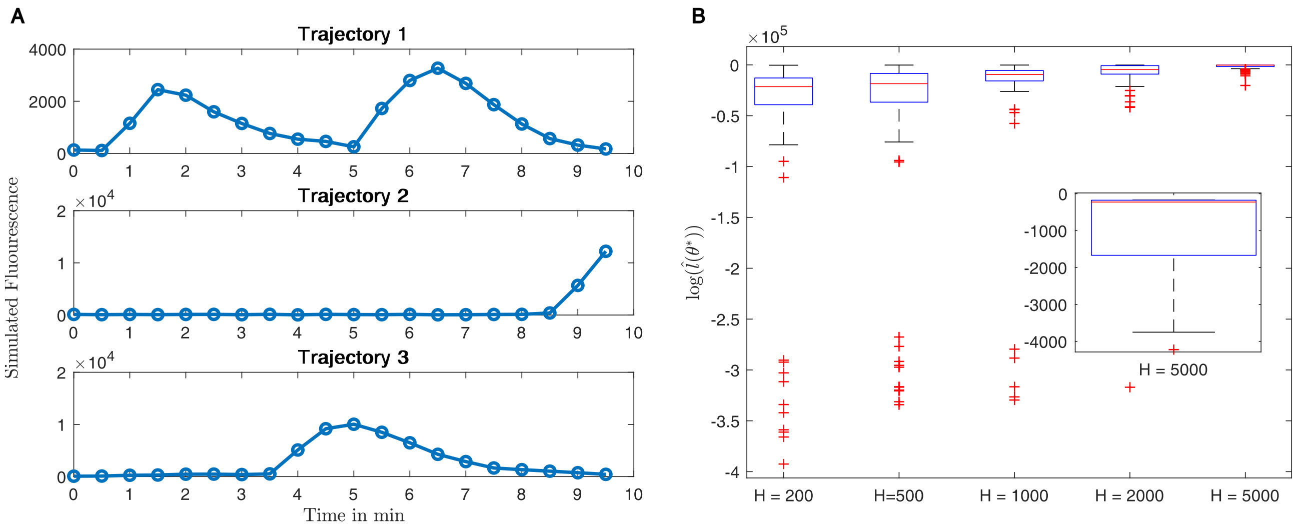

**Figure S4:** **A:** 3 example trajectories of simulated Lac-Gfp data, measured at 29 time points. **B:** Log likelihood approximation for the real parameter $\theta^*$ for the first trajectory with different number of particles $H$ for the particle filter.

**Table S3:** Prior distributions of the parameters for the transcription model.

| Parameter | Meaning | Prior interval |
|---|---|---|
| $k_{\text{on}}$ | promoter binding rate | $[10^{-5}, 100]$ |
| $k_{\text{off}}$ | promoter unbinding rate | $[10^{-5}, 100]$ |
| $k_{\text{r}}$ | mRNA transcprition | $[0.1, 500]$ |
| $\lambda$ | transcription rate | $[2, 8]$ |

For the transcription model inference problem presented in this paper, each parameter was assigned an independent uniform log prior distribution in the interval listed in the table.

1. $g_{\text{off}} \xrightarrow{k_{\text{on}}} g_{\text{on}}$ Promoter binding to gene.
2. $g_{\text{on}} \xrightarrow{k_{\text{off}}} g_{\text{off}}$ Promoter unbinding from gene.
3. $g_{\text{on}} \xrightarrow{k_r} g_{\text{on}} + mRNA_1$ Transcription process
4. $RNA_1 \xrightarrow{\lambda} RNA_2$ Transcription process
5. $RNA_2 \xrightarrow{\lambda} RNA_3$ Transcription process
6. $RNA_3 \xrightarrow{\lambda} RNA_4$ Transcription process
7. $RNA_4 \xrightarrow{\lambda} RNA_5$ Transcription process
8. $RNA_5 \xrightarrow{\lambda} RNA_6$ Transcription process
9. $RNA_6 \xrightarrow{\lambda} RNA_7$ Transcription process
10. $RNA_7 \xrightarrow{\lambda} RNA_8$ Transcription process
10. $RNA_8 \xrightarrow{\lambda} \emptyset$ Transcription process

The parameters are shown in table S3. Note that the expected total transcription time is $\frac{8}{\lambda}$, this is why the prior for $\lambda$ was chosen between 2 and 8.

The measurements are assumed to be the noisy read out of the total number of $mRNA$, where each $mRNA$ molecule emits fluourescence with mean $\mu = 1$ and variance $\sigma^2 = 1/8$. We also assume a background fluourescence of $\mathcal{B} = \mathcal{N}(0, 4)$. The final measurement thus reads

$$y(t) = \mathcal{N}\left(\mu \sum_{i=1}^{7} mRNA_i, \sigma^2 \sum_{i=1}^{7} mRNA_i\right) + \mathcal{B}.$$
